# Single cell approaches define forebrain neural stem cell niches and identify microglial ligands that enhance precursor-mediated remyelination

**DOI:** 10.1101/2024.03.22.586277

**Authors:** Ashleigh Willis, Danielle Jeong, Yunlong Liu, Marissa A. Lithopoulos, Scott Yuzwa, Paul W. Frankland, David R. Kaplan, Freda D. Miller

## Abstract

Here we used single cell RNA-sequencing and single cell spatial transcriptomics to characterize the forebrain neural stem cell (NSC) niche under homeostatic and injury conditions. We define the dorsal and lateral ventricular-subventricular zones (V-SVZ) as two distinct neighborhoods, and show that following white matter injury, dorsal NSCs are locally activated to make oligodendrocytes for remyelination. This activation is coincident with a robust increase in transcriptionally-distinct microglia in the dorsal V-SVZ niche. We modeled ligand-receptor interactions within this changing niche and identified two remyelination-associated microglial ligands, IGF1 and OSM, that promote precursor proliferation and oligodendrogenesis in culture. Infusion of either ligand into the lateral ventricles also enhanced oligodendrogenesis, even in the lateral V-SVZ, where NSCs normally make neuroblasts. These data support a model where gliogenesis versus neurogenesis is determined by the local NSC neighborhood and where injury-induced niche alterations promote NSC activation, local oligodendrogenesis, and likely contribute to myelin repair.

## INTRODUCTION

Recruitment of endogenous neural precursor cells (NPCs) to promote brain repair is one idea gaining increased traction based in part on early clinical data with drugs that regulate NPCs such as metformin^1,2^. Such a strategy is of particular interest for myelin repair, since newly-generated oligodendrocytes are capable of functionally integrating into adult white matter tracts (reviewed in ^3^). Moreover, myelin deficits and abnormal white matter indices are observed in many pathological neural conditions ranging from multiple sclerosis to neurodevelopmental disorders and traumatic brain injury (reviewed in ^4^), and thus any strategy that effectively increased oligodendrogenesis would be of considerable clinical importance. Most work studying this possibility focuses on oligodendrocyte precursor cells (OPCs), which are broadly distributed throughout the brain. Indeed, differentiation of parenchymal OPCs into oligodendrocytes is known to occur during spontaneous remyelination following a demyelinating injury^5,6^. However, in humans OPCs are unable to completely restore lost myelin particularly with age and progressive disease^7^, highlighting the need to consider additional endogenous sources for generating myelinating oligodendrocytes^8^.

A second potential endogenous oligodendrocyte source is forebrain NSCs. During developmental myelination, V-SVZ NSCs produce the large majority of corpus callosum oligodendrocytes and forebrain parenchymal OPCs^9^. This NSC-mediated oligodendrogenesis continues to some extent throughout life^10^, particularly in the dorsal V-SVZ, which is adjacent to the corpus callosum and is thought to be biased towards gliogenesis (reviewed in ^11–13^). This is in contrast to NSCs in the lateral V-SVZ, which preferentially generate olfactory bulb interneurons. In rodents, V-SVZ NSCs also contribute oligodendrocytes to white matter tracts following experimental demyelinating injury^14–18^ and even in humans there is an apparent increase in NSC-mediated oligodendrogenesis in post-mortem brain tissue from patients with long-term demyelinating disorders^19^.

How then might we recruit endogenous NSCs to promote remyelination? While the aforementioned studies have established that NSCs can contribute to remyelination, we don’t yet understand the cellular state of NSCs following white matter injury, nor do we understand the NSC niche in homeostatic versus injury conditions. Here, we have chosen to address these questions using a combination of lineage tracing, single cell RNA-sequencing (scRNA-seq) and single cell Xenium and MERFISH-based spatial transcriptomics. We have used these approaches to study the adult murine V-SVZ under homeostatic conditions and during injury-induced remyelination. These analyses have enabled us to define distinct V-SVZ neighborhood and ligand environments and to identify two ligands, IGF1 and OSM, that are locally increased in microglia in the dorsal V-SVZ during remyelination and enhance NPC-mediated oligodendrogenesis. Based on these findings, we propose a model where differing dorsal and lateral V-SVZ neighborhoods normally regulate neurogenesis versus gliogenesis and that following white matter injury, microglia alter the dorsal NSC niche, in part by secreting IGF1 and OSM, to locally increase oligodendrogenesis and promote remyelination. These findings have implications for our understanding of the NSC niche, of the role that microglia play in demyelinating disorders, and ultimately, for the development of endogenous remyelination strategies.

## RESULTS

### Nestin-expressing NPCs in the adult V-SVZ contribute to ongoing oligodendrogenesis and this increases two- to three-fold during remyelination

Adult V-SVZ NSCs of the dorsal wall have gliogenic potential throughout life^10,20^. To better-understand their baseline contribution to adult corpus callosum oligodendrogenesis, we performed lineage tracing using a well-characterized *Nestin^CreERT^*^2^ mouse line^80^ crossed to mice carrying a floxed *Rosa26R-EYFP* reporter allele (*Nestin^CreERT^*^2^*^/+;^R26R-EYFP*; called *NestinCreErt2-Eyfp* mice from hereon). When these crossed mice are exposed to tamoxifen, *Nestin*-expressing NSCs, transit-amplifying precursor cells (TAPs) and their progeny are tagged with EYFP. *Nestin*-expressing ependymal cells will also be labelled. Since it is still not clear whether ependymal cells function as neural precursors following injury^21–23^, and since TAPs will also be labelled, we have specified that this strategy will label NPCs, and not specifically NSCs.

We analyzed the forebrain of these mice after five consecutive days of tamoxifen treatment. As predicted, only cells around the V-SVZ were EYFP-positive (Suppl. Fig. 1A). We then waited for 8 weeks post-tamoxifen, and analyzed NPC progeny by immunostaining forebrain sections through the V-SVZ and corpus callosum, identifying oligodendrocyte precursor cells (OPCs) by their expression of OLIG2 and PDGFRα and oligodendrocytes by their expression of OLIG2 and CC1 (Suppl. Fig. 1B). OPCs and oligodendrocytes were observed throughout the corpus callosum as expected. Quantification showed that about 8% of OPCs and 11-14% of oligodendrocytes in both the rostral and caudal corpus callosum were EYFP-positive (Fig. 1A) indicating they had been generated in 8 weeks following tamoxifen administration.

**Figure 1.**
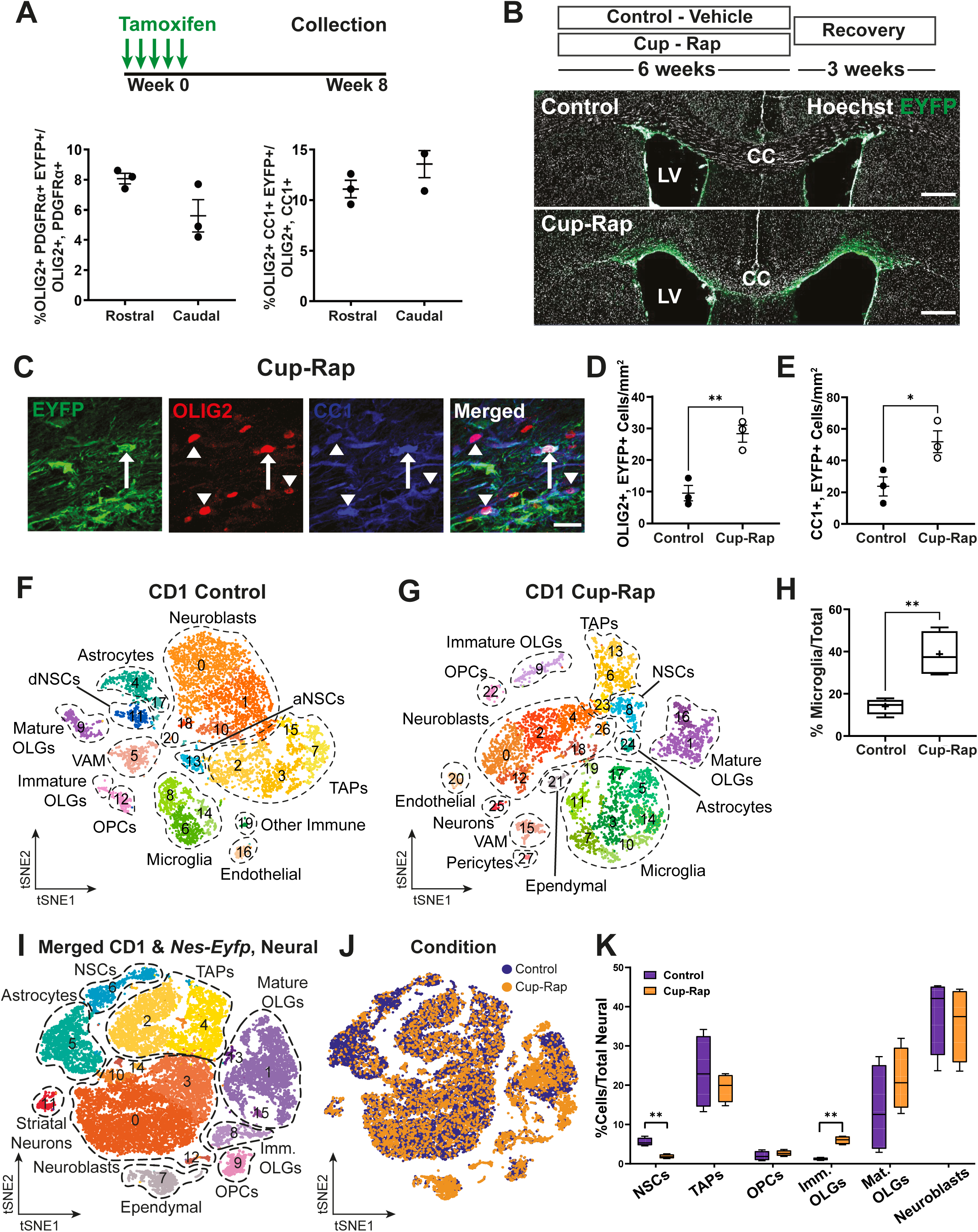
Analysis of the adult V-SVZ by lineage tracing and scRNA-seq identifies increased microglia and NPC-mediated oligodendrogenesis during remyelination. Also see Suppl. Figures 1 and 2. (A) Eight-week-old *NestinCreErt2-Eyfp* mice were injected with tamoxifen and coronal forebrain sections at the level of the corpus callosum and lateral ventricles were analyzed eight weeks later by immunostaining for EYFP, OLIG2, and PDGFRα or CC1. The top panel shows a schematic of the experiment. The left scatter plot shows quantification of the rostral and caudal corpus callosum for percentages of EYPF-positive, OLIG2-positive, PDGFRα-positive OPCs and the right plot percentages of EYFP-positive, OLIG2-positive, CC1-positive oligodendrocytes. Each dot denotes a different mouse. **(B)** Six-week-old *NestinCreErt2-Eyfp* mice were treated with cuprizone-rapamycin or vehicle for six weeks, and analyzed three weeks-post-treatment as in the schematic. Shown are representative low-magnification micrographs of coronal sections of the V-SVZ and corpus callosum (CC), analyzed for EYFP (green) and counterstained with Hoechst 33258 (white). LV = lateral ventricle. Scale bars = 300 µm. **(C)** Representative high-magnification confocal images of the corpus callosum of a cuprizone-rapamycin-treated *NestinCreErt2-Eyfp* mouse three weeks post-treatment, analyzed for EYFP and for OLIG2 and CC1 immunoreactivity. The arrow denotes a triple-labeled NPC-derived mature oligodendrocyte (arrow), and arrowheads EYFP-negative, OLIG2-positive, CC1-positive resident mature oligodendrocytes. Scale bars = 20 µm. **(D, E)** Quantification of sections as in (C) for the relative densities of (D) EYFP-positive, OLIG2-positive or (E) EYFP-positive, CC1-positive cells in the rostral corpus callosum. Each dot represents an individual animal. *p<0.05, **p<0.01. **(F, G)** Adult CD1 mouse dorsal and lateral V-SVZ tissue was collected from control vehicle (F) or cuprizone-rapamycin treated (G) mice and scRNA-seq was performed. Shown are t-SNE cluster map visualizations of all V-SVZ transcriptomes from the merger of two runs each of the two experimental groups (individual datasets are shown in Suppl. Fig. 1C-H). For the remyelinating dataset, both runs were 3 weeks post-treatment, and for the controls, one was at 3 weeks and the other immediately after treatment. Transcriptionally-distinct clusters are colour-coded, and the plot is annotated for the cell types within these clusters, as determined by analysis of well-characterized marker genes. dNSC = dormant NSCs, aNSC = activated NSCs, NB = neuroblasts, OL = oligodendrocytes. VAM = vasculature-associated mesenchymal cells. **(H)** Box plots showing percentages of microglia relative to total cells in the individual scRNA-seq datasets (Suppl. Fig. 1C-K; Suppl. Fig. 2A-C). N = 4 independent scRNA-seq datasets for each condition. **p<0.01. **(I, J)** Neural cell transcriptomes from the CD1 and *NestinCreErt2-Eyfp* control and cuprizone-rapamycin-treated datasets were subsetted, merged, and subjected to one round of Harmony-mediated batch correction (4 control datasets, shown in Suppl. Fig. 1C, D, I, J and 4 cuprizone-rapamycin-treated datasets, shown in Suppl. Fig. 1F, G and Suppl. Fig. 2A, B). Panel (I) shows the cluster t-SNE, where transcriptionally distinct clusters are numbered and annotated for neural cell types as determined by gene expression profiles (Suppl. Fig. 2E, F), and (J) shows the experimental condition for each transcriptome. Abbreviations are as in panel F. **(K)** Box plots showing the percentage of total neural cells that were NSCs, TAPs, olfactory neuroblasts, or cells of the oligodendrogenic lineage in individual control and cuprizone-rapamycin treated datasets. N = 4 independent scRNA-seq datasets for each condition. **p<0.01.

We asked if this adult NPC-mediated oligodendrogenesis was altered during injury-induced remyelination. We used a well-established model^24^ where six weeks of cuprizone-rapamycin treatment causes widespread demyelination. Once this treatment is withdrawn, the corpus callosum is completely remyelinated in eight weeks. We confirmed corpus callosum demyelination by immunostaining for myelin basic protein (MBP) 1.5 weeks post-treatment (data not shown). We then treated *NestinCreErt2-Eyfp* mice with cuprizone-rapamycin for six weeks, administered tamoxifen for five days to label *Nestin*-expressing NPCs, and analyzed the forebrain three weeks post-treatment, at the midpoint of remyelination. As controls, mice were fed the same chow without cuprizone and injected with vehicle rather than rapamycin. Immunostaining revealed a robust increase in EYFP-positive cells in the dorsal V-SVZ and corpus callosum in the remyelinating brains (Fig. 1B). Quantification of triple-labelled sections revealed significant 2-to-3-fold increases in EYFP-positive, CC1-positive oligodendrocytes and EYFP-positive, OLIG2-positive, CC1-negative immature oligodendrocytes/OPCs (Fig. 1C-E). Thus, remyelination is accompanied by increased NPC-mediated oligodendrogenesis, consistent with previous reports^14–18^.

### Single cell transcriptomic analysis identifies increases in microglia and NPC-derived immature oligodendrocytes during remyelination

To obtain a global overview of the NSC niche and to understand how it might regulate oligodendrogenesis, we used scRNA-seq to characterize the control and three-week remyelinating V-SVZ. Specifically, we treated CD1 wildtype mice with cuprizone-rapamycin for six weeks, withdrew the treatment, and three weeks later collected the lateral and dorsal V-SVZ and adjacent corpus callosum. As controls, we analyzed tissue from mice fed normal powdered chow and injected with vehicle for six weeks, collected immediately or three weeks post-vehicle; similar results were obtained at both timepoints. We also isolated tissue from *NestinCreErt2-Eyfp* mice injected with tamoxifen for 5 days prior to the same treatment in order to combine scRNA-seq with lineage-tracing. In all cases we used fluorescence-activated cell sorting to enrich for viable cells, and sequenced single cells using the 10X Genomics platform. We analyzed two runs for each group (8 runs total), with at least 4 mice per run.

To analyze the resultant transcriptomes we used our previously described pipeline^25–28^. After filtering the datasets for low quality cells with for example few expressed genes or high mitochondrial proportions and cell doublets, we obtained 2573 to 11030 single cell transcriptomes in each of the eight independent datasets (39,905 cells total). Genes with high variance were used to compute principal components as inputs for projecting cells in two-dimensions using t-distributed stochastic neighbor embedding (t-SNE) and clustering performed using a shared nearest neighbors-cliq (SNN-cliq)-inspired approach built into the Seurat R package at a range of resolutions. We used gene expression overlays for well-validated marker genes to define cell types (Suppl. Fig. 1C-K; Suppl. Fig. 2A-C).

This analysis identified the expected V-SVZ and corpus callosum cell types in all datasets, including NSCs, TAPs, neuroblasts, astrocytes, OPCs, immature and mature oligodendrocytes, ependymal cells, microglia, vasculature-associated mesenchymal cells and endothelial cells (Fig. 1F, G; Suppl. Fig. 1C-K; Suppl. Fig. 2A-C). However, relative cellular proportions differed in control versus cuprizone-rapamycin-treated datasets with the most evident being a significant two-to-three-fold increase in microglial numbers during remyelination, as quantified by calculating microglial cell percentages in each individual dataset (Fig. 1H).

We subsetted, merged and reanalyzed neural cell transcriptomes from all the datasets to directly compare control and remyelinating cells. We corrected for variability between the CD1 and *NestinCreErt2-Eyfp* mouse datasets using one round of Harmony-mediated batch-correction^29^ (Suppl. Fig. 2D). t-SNE-based clustering and gene expression overlays (Fig. 1I; Suppl. Fig. 2E, F) identified all expected cell types in the merged dataset. Analysis of the dataset of origin (Fig. 1J) showed that control and remyelinating transcriptomes were largely intermingled with two exceptions, microglia (discussed further below) and the oligodendroglial lineage. Specifically, the immature oligodendrocyte cluster was almost completely comprised of remyelinating cells, and the control versus remyelinating mature oligodendrocytes were distinctly clustered, presumably reflecting different states of maturation. Quantification of relative cell proportions in individual datasets (Fig. 1K) confirmed a significant increase in total immature oligodendrocytes during remyelination and a coincident significant decrease in total NSCs. By contrast, the relative proportions of TAPs and neuroblasts were not significantly altered.

We asked if NPCs contributed to this increased oligodendrogenesis by analyzing the *NestinCreErt2-Eyfp* datasets (Fig. 2A, C). Specifically, we subsetted neural cells from the total individual datasets (Suppl. Fig. 1I-K; Suppl. Fig. 2A-C), and merged and reanalyzed transcriptomes from the two control versus two cuprizone-rapamycin datasets. Quantification of *Eyfp* mRNA-positive cells in these merged datasets (Fig. 2B, D and E) showed the NPC labeling strategy was efficacious; 60-70% of NSCs and almost 90% of TAPs were *Eyfp*-positive in both conditions. Analysis of downstream NPC progeny showed that about 60% of neuroblasts were labelled in the control datasets, and this was unchanged during remyelination. However, the oligodendrocyte lineage differed between the two conditions. In controls, only 5-7% of OPCs, immature oligodendrocytes and mature oligodendrocytes were *Eyfp*-positive, but this labeling index was increased 3 to 6-fold during remyelination (Fig. 2E). Thus, in agreement with the morphological data (Fig. 1A-E), V-SVZ NPCs increased their genesis of oligodendroglial cells during injury-induced remyelination.

**Figure 2.**
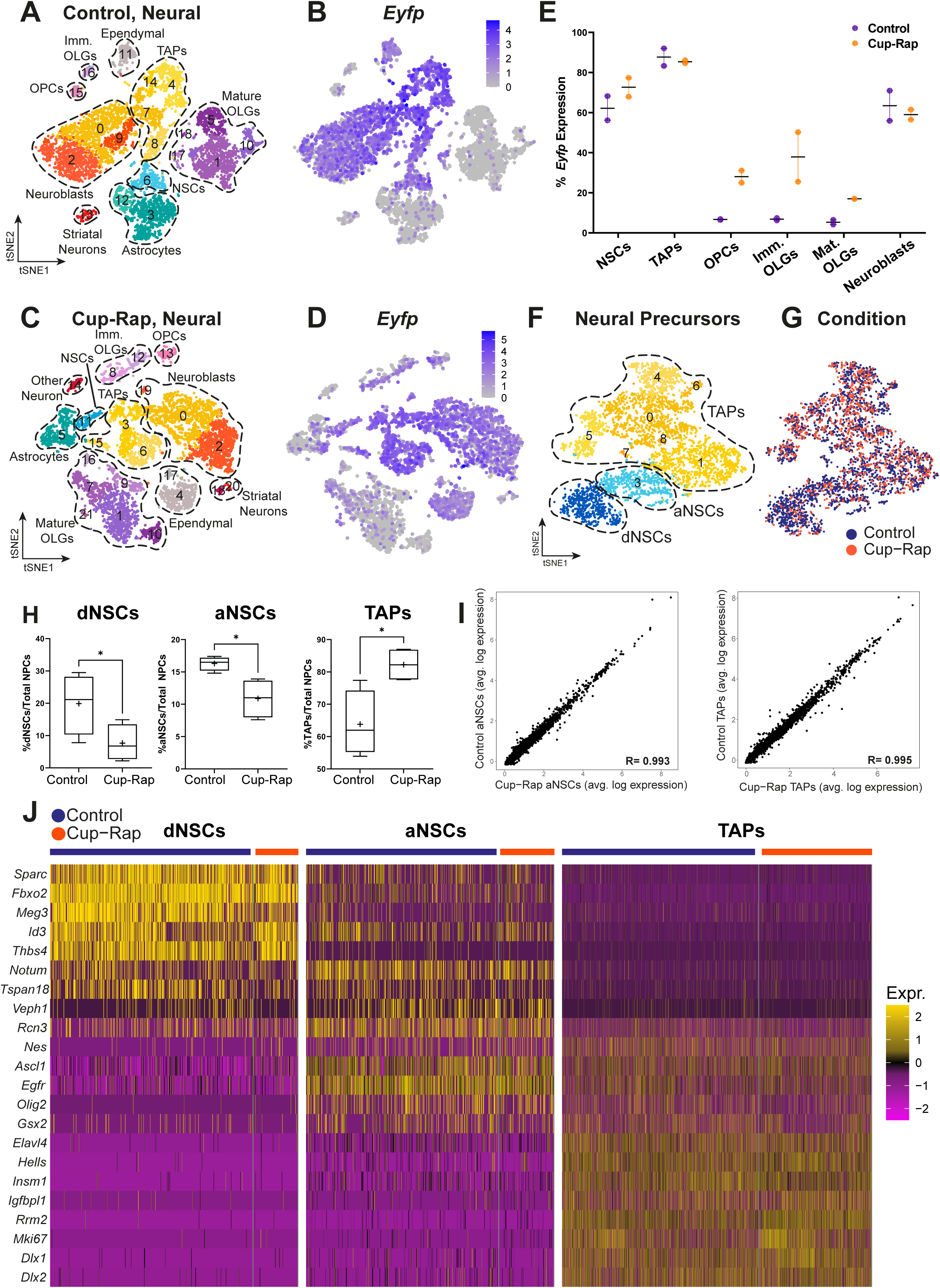
V-SVZ NSCs are activated and increase their genesis of oligodendroglial lineage cells during injury-induced remyelination. Also see Suppl. Figures 1 and 2. (A-D) t-SNE visualizations of subsetted, merged adult V-SVZ neural cells from the three week post-treatment *NestinCreErt2-Eyfp* control (A, B) and cuprizone-rapamycin (C, D) datasets (shown in Suppl. Fig. 1I-K and Suppl. Fig. 2A-C). In (A and C) transcriptionally distinct clusters are numbered and annotated for cell types. In (B and D), the cluster t-SNE is overlaid for expression of *Eyfp*, with expression levels coded as per the adjacent keys. **(E)** Scatter plot showing the percentage of selected cell types relative to total neural cells from the individual control and cuprizone-rapamycin treated *NestinCreErt2-Eyfp* datasets (4 datasets total). Each dot represents values from one dataset. **(F, G)** t-SNE visualization of merged control and cuprizone-rapamycin treated adult V-SVZ NPCs (NSCs and TAPs) subsetted from the merged neural cell dataset shown in Fig. 1I, with one round of Harmony-mediated batch correction. (F) shows the cluster t-SNE, with the three NPC types annotated based on their gene expression profiles. (G) shows the same t-SNE cluster visualization with transcriptomes colour-coded for treatment condition. **(H)** Box plots showing the percentage of dNSCs, aNSCs and TAPs relative to total NPCs in each of the 8 individual scRNA-seq datasets (Suppl. Fig. 1C-K; Suppl. Fig. 2A-C). N = 4 per condition. *p<0.05. **(I)** Pearson correlation analysis of each detected gene in aNSCs (left) and TAPs (right) from the dataset shown in (F), comparing averaged gene expression of control (y-axis) and cuprizone-rapamycin treated (x-axis) transcriptomes. **(J)** Single cell heatmap showing expression of select mRNAs that distinguish dNSCs, aNSCs and TAPs, comparing their expression in individual control versus cuprizone-rapamycin treated transcriptomes from the dataset shown in (F and G). Each row corresponds to a gene and each column line represents the expression level for that particular mRNA in a single cell. Gene expression levels are coded as per the adjacent key.

### Demyelination causes activation of V-SVZ NSCs and TAPs but does not alter their transcriptional identities

To better-understand the observation that NSCs are proportionately decreased during remyelination (Fig. 1K), we subsetted and reanalyzed NSCs and TAPs from the neural cell dataset (clusters 2, 4 and 6 from Fig. 1I). We defined dormant and activated NSCs and TAPs using previously published marker genes (Fig. 2F; Suppl. Fig. 2G, H; also see the heatmap in Fig. 2J). Analysis of the dataset of origin (Fig. 2G) showed that control and remyelinating NSCs and TAPs were intermingled (Fig. 2H), but quantification of individual datasets demonstrated that their relative proportions were significantly different (Fig. 2H). In controls 22% of NPCs were dormant NSCs, 17% activated NSCs, and the remaining 63% TAPs. During remyelination dormant and activated NSCs were decreased two-fold and 50% respectively, while TAPs were increased by 25%. However, the proliferative index was similar under both conditions; Cyclone^30^ predicted that only TAPs were proliferative and that similar proportions were cycling in both conditions (a mean of 42% +/- 1.9 SEM and 45% +/- 4.5 SEM in control and cuprizone-rapamycin, respectively).

These data suggest that following demyelination, dormant NSCs are activated to generate TAPs and ultimately their downstream oligodendroglial progeny. We asked if NPC transcriptional states were also altered using Pearson correlation of average gene expression with the uncorrected data. Dormant NSCs, activated NSCs and TAPs were highly similar in control versus remyelinating conditions, with r values of 0.97 to 0.99 (Fig. 2I, Suppl. Fig. 2I). These similarities were also apparent in a single cell heatmap of genes enriched in each of the three NPC populations (Fig. 2J). These included genes enriched specifically in dormant NSCs (*Thbs4, Id3, Sparc, Fbxo2* and *Meg3*), in both dormant and activated NSCs (*Notum*, *Tspan18* and *Veph1*), in both activated NSCs and TAPs (*Egfr*, *Ascl1*, *Gsx2,* and *Olig2*) and specifically in TAPs (*Hells, Rrm2, Insm1, Igfbpl1, Elavl4, Dlx1, Dlx2* and *Ki67*). Thus, V-SVZ NPCs retain their control transcriptional state during remyelination.

### Remyelination-associated microglia display an altered transcriptional state

These data suggest that following demyelination the NSC niche is altered so that NSCs are activated and oligodendrogenesis is increased. Since an increase in microglia was the largest cellular alteration we observed, we characterized these cells in more detail by subsetting them from a merge of all the CD1 mouse transcriptomes (clusters 3, 11-14 and 22 from Suppl.Fig. 3A, B). Cluster visualization and gene expression overlays (Fig. 3A-C; Suppl. Fig. 3C, D) showed that one group of clusters was almost completely comprised of microglia from the remyelinating condition. We have termed these remyelination-enriched microglia from hereon. A second group was largely, but not exclusively, comprised of microglia from the control condition. We have termed these homeostatic microglia recognizing that this group also includes some microglia from the remyelinating condition and that these cells likely display dynamic and diverse functional and morphological states.

**Figure 3.**
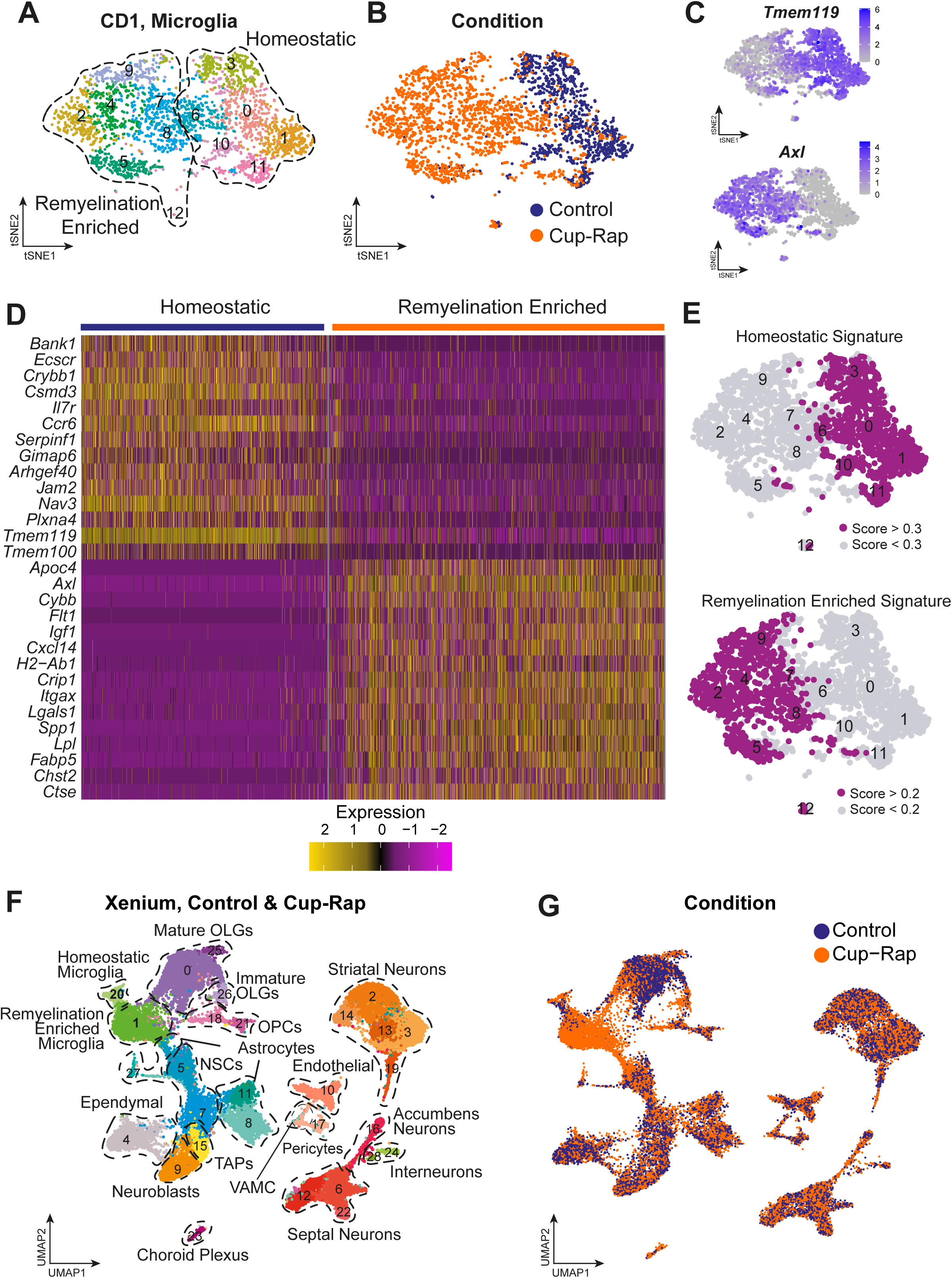
Microglia display an altered transcriptional state during injury-induced remyelination. Also see Suppl. Figure 3 and Suppl. Tables 1 and 2. (A-C) t-SNE visualizations of merged control and cuprizone-rapamycin treated adult microglia subsetted from the merged total CD1 datasets (two control and two cuprizone-rapamycin treated; shown in Suppl. Fig. 3A, B). (A) shows the cluster t-SNE, where the two groups of transcriptionally-distinct microglia are outlined. (B and C) show the same cluster visualization where each transcriptome is colour-coded for treatment condition (B) or is overlaid for expression of *Tmem119* and *Axl*, mRNAs enriched in the two microglial groups (C). Expression levels are coded as per the adjacent keys. **(D)** Single cell heatmap showing expression of select mRNAs differentially expressed in the two microglial groups shown in (A). Each row corresponds to a gene and each column line represents the expression level for that particular mRNA in a single cell. Expression levels are coded as per the adjacent key. **(E)** t-SNE visualization as in (A) overlaid for gene expression signatures comprised of 40 and 65 significantly-increased mRNAs in the homeostatic versus remyelination-enriched microglia, respectively (see Methods for gene names). Red denotes cells with scores > 0.2 or > 0.3, as specified. **(F, G)** Coronal sections from vehicle or cuprizone-rapamycin treated mice were obtained 3 weeks post-treatment and analyzed by Xenium-based single cell multiplexed *in situ* hybridization with a probeset targeting 347 genes (see Suppl. Table 2). The ROI was defined as the region around the V-SVZ plus the corpus callosum (see Fig. 4A). Cellular transcriptomes within the ROI of an individual section were analyzed after removal of poor-quality transcriptomes. Datasets from different sections and conditions were then merged and cell types identified by marker gene expression. (F) shows UMAP cluster visualization of the merged transcriptomes, annotated for cell types identified using well-characterized marker genes (see Suppl. Fig. 3E-H) while (G) shows the same cluster UMAP where transcriptomes are color-coded for treatment condition. Each dot represents a single transcriptome. VAMC = vasculature-associated mesenchymal cell, OLGs = oligodendrocytes.

Differential gene expression analysis (Suppl. Table 1) showed that these two microglial groups were distinguished by 1386 genes that were expressed in at least 10% of homeostatic or remyelination enriched microglia (average fold change > 2, Bonferroni adjusted p-value < 0.05). The homeostatic microglia were enriched for 486 genes previously associated with unperturbed adult microglia such as *Tmem119, Sall1* and *P2ry12,* as well as other genes including *Crybb1, Il7r* and *Ecscr*. The remyelination-enriched microglia were enriched for microglial genes previously associated with the injured/degenerating/inflamed brain such as *Axl* and *Cst7,* as well as genes like *Apoc4, Cybb* and *Flt1* (900 genes total) (Fig. 3C, D; Suppl. Fig. 3C, D). Notably, the remyelination-enriched microglia also displayed increased expression of several ligand mRNAs, including *Igf1, Osm* and *Cxcl14* (Suppl. Table 1; Fig. 3D). We used these differentially-expressed genes to generate transcriptional signatures for the two microglial transcriptional states (Fig. 3E).

### Single cell spatial transcriptomics define distinct NSC niches in the control and remyelinating V-SVZ

One limitation of scRNA-seq data is that it does not provide spatial information. We therefore performed *in situ* hybridization-based single cell spatial transcriptomics with the Xenium platform^31^, analyzing control and remyelinating forebrain sections. We used a custom 347 gene probeset (Suppl. Table 2) that allowed identification of the relevant V-SVZ cell types, as informed by the scRNA-seq data. We analyzed a region of interest (ROI) that included the corpus callosum and V-SVZ plus adjacent cells (for example, see Fig. 4A). We removed a small number of objects with low transcript counts (likely cellular fragments), and merged cells from different sections/animals from the same condition. Uniform manifold approximation and projections (UMAPs) (Suppl. Fig. 3E, F) showed that for both conditions transcriptomes from different sections were well-integrated, and marker gene analysis identified the expected cell types (Suppl. Fig. 3E, F). We then merged the uninjured and remyelinating datasets to allow a direct comparison of cells and neighborhoods in these two conditions (Fig. 3F, G; Suppl. Fig. 3G, H). We ensured correct identification of NSCs versus astrocytes within the merged dataset by subsetting and analyzing NSCs, TAPs, astrocytes and ependymal cells (Suppl. Fig. 4A, B), utilizing previously-identified marker genes^26,27^ (see Experimental Methods).

**Figure 4.**
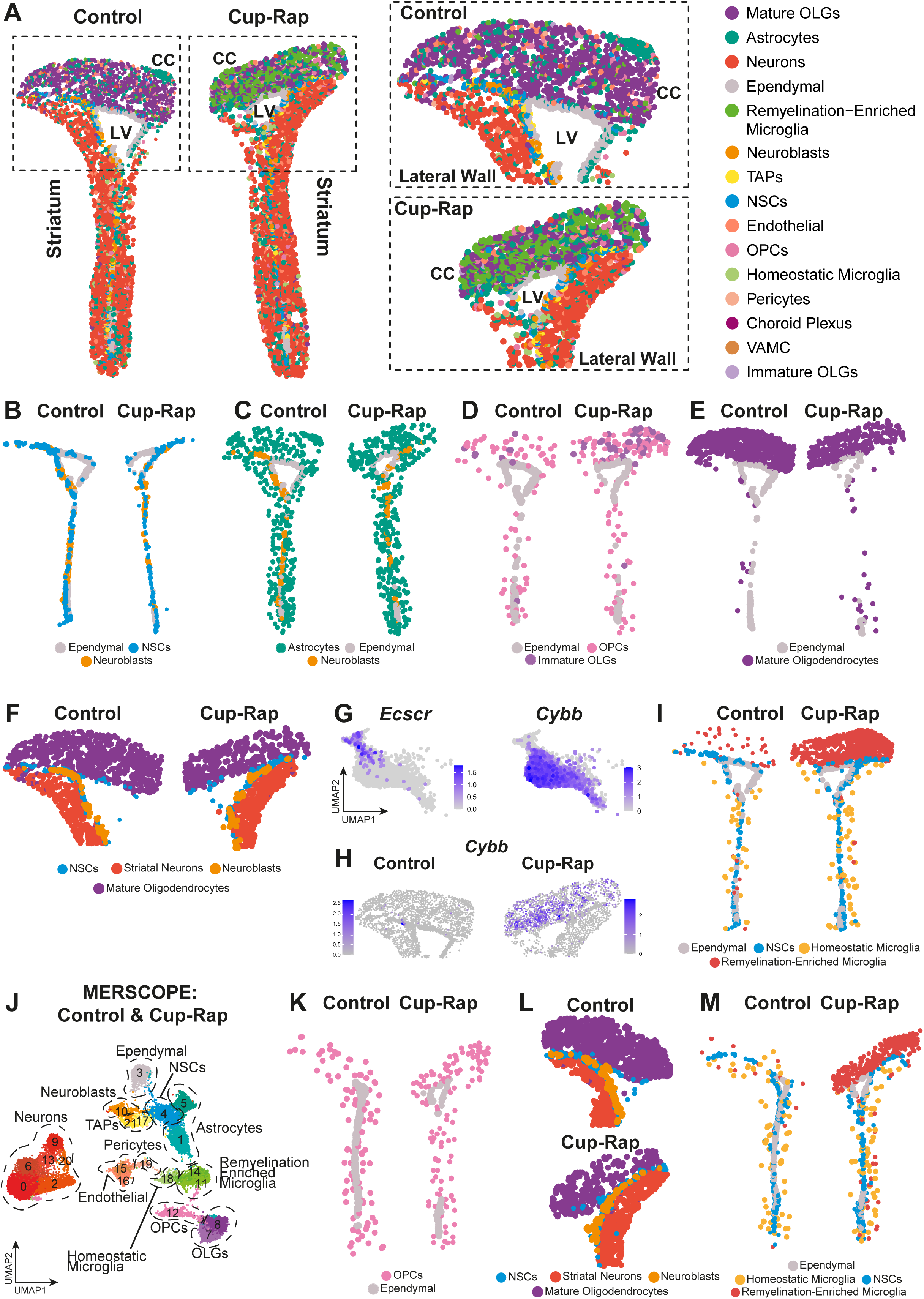
Xenium and Merfish-based single cell transcriptomics define distinct dorsal and lateral wall NSC niches in the control and remyelinating V-SVZ. Also see Suppl. Figure 4 and Suppl. Table 2. (A-I) Spatial and UMAP cluster plots of the cell types identified in the annotated Xenium-based analysis (the UMAPs in Figure 2F, G) showing the region around the lateral ventricles and the corpus callosum (the ROI). All spatial plots (A-F, H and I) show one representative section each for the control and remyelinating (Cup-Rap) conditions. **(A)** Spatial plot of the entire ROI, showing all of the cell types, color-coded to the right. The boxed region including the dorsal V-SVZ, corpus callosum and top of the lateral V-SVZ are shown in an enlarged view to the right. Each dot represents an individual cell. VAMC = vasculature-associated mesenchymal cells, OLGs = oligodendrocytes, LV = lateral ventricle, CC = corpus callosum. **(B)** Spatial plots of NSCs (blue), neuroblasts (orange) and ependymal cells (grey) in the ROI. **(C)** Spatial plots of astrocytes (green), neuroblasts (orange) and ependymal cells (grey) in the ROI. **(D)** Spatial plots of OPCs (pink), immature oligodendrocytes (purple) and ependymal cells (grey) in the ROI. **(E)** Spatial plots of mature oligodendrocytes (dark purple) and ependymal cells (grey) in the ROI. **(F)** Spatial plots showing differences in the dorsal and lateral V-SVZ neighborhoods, with striatal neurons colored red, neuroblasts orange, NSCs blue and mature oligodendrocytes dark purple. The plot includes the dorsal V-SVZ, the corpus callosum and the top of the lateral V-SVZ. **(G)** Gene expression overlays for *Ecscr* and *Cybb*, mRNAs differentially expressed in homeostatic versus remyelination-enriched microglia, shown on the enlarged microglial clusters (1 and 20) from the annotated UMAP in Figure 3F. Relative expression levels are coded as per the adjacent keys. **(H)** Spatial plot for *Cybb* mRNA expression in the dorsal V-SVZ, corpus callosum and top part of the lateral V-SVZ of representative control and remyelinating sections. Relative expression levels are coded as per the adjacent keys. **(I)** Spatial plots showing homeostatic microglia (yellow), remyelination-enriched microglia (red), NSCs (blue) and ependymal cells (grey) in the ROI. **(J-M)** Coronal sections from vehicle or cuprizone-rapamycin treated mice were obtained 3 weeks post-treatment and analyzed by MERSCOPE-based single cell multiplexed *in situ* hybridization with a probeset targeting 300 genes (in Suppl. Table 2). The same ROI was analyzed as for the Xenium analysis. Cellular transcriptomes from individual sections were analyzed, datasets from different sections and conditions were merged and cell types identified by marker gene expression. **(J)** UMAP cluster visualization of the merged control and cuprizone-rapamycin treated transcriptomes, annotated for cell types as identified by marker gene expression (Suppl. Fig. 4E-H). Each dot represents a single transcriptome. OLGs = oligodendrocytes. **(K-M)** Spatial plots of the cell types identified in the annotated MERSCOPE-based analysis in (J) showing the V-SVZ and corpus callosum (the ROI). All plots show one representative section each for the control and remyelinating (Cup-Rap) conditions and each dot is a cell. **(K)** shows OPCs (pink) and ependymal cells (grey). **(L)** is an expanded view of the dorsal V-SVZ, corpus callosum and top of the lateral V-SVZ showing striatal neurons (red), neuroblasts (orange), NSCs (blue) and mature oligodendrocytes (dark purple). **(M)** shows homeostatic microglia (yellow), remyelination-enriched microglia (red), NSCs (blue) and ependymal cells (grey).

Spatial plots (Fig. 4A-F; Suppl. Fig. 4C) showed that in the control V-SVZ cells were in their expected locations, with ependymal cells lining the ventricles, OPCs and astrocytes scattered throughout the grey and white matter, and oligodendrocytes enriched within the corpus callosum. Vasculature-associated cells (endothelial cells and pericytes) were located throughout the tissue, and the different neuronal populations were appropriately localized in the parenchyma. As predicted, NSCs were closely apposed to ependymal cells throughout the V-SVZ (Fig. 4B). However, their other cellular neighbors varied (Fig. 4A, F). In particular, NSCs in the dorsal V-SVZ, which are biased to make glia^20,32^, were closely-associated with oligodendrocytes. By contrast, NSCs within the lateral V-SVZ, which predominantly generate olfactory bulb interneurons^33^, were instead closely-associated with their neuroblast progeny and adjacent striatal neurons. Thus, different V-SVZ NSC cellular neighborhoods correlate with distinct profiles of cell genesis.

We next compared the dorsal and lateral V-SVZ neighborhoods in the control versus remyelinating conditions. Analysis of the dataset of origin for the merged Xenium dataset (Fig. 3F, G) showed that with two exceptions (discussed below) most cells from the two conditions were co-clustered, consistent with the scRNA-seq-based conclusion that their transcriptional states were largely unaltered during remyelination. Spatial plots of the remyelinating cells (Fig. 4A-F) showed that during remyelination ependymal cells lined the ventricles while OPCs, astrocytes and vasculature-associated cells were scattered throughout the grey and white matter, as seen in controls. Moreover, the organization of the lateral V-SVZ niche was largely unaffected, with neuroblasts, NSCs and neurons localized as they were in the control situation. By contrast, the dorsal V-SVZ neighborhood was altered during remyelination. The first major difference involved the oligodendroglial lineage (Fig. 3F, G); during remyelination there were relatively more immature oligodendrocytes and the mature oligodendrocytes differed transcriptionally from controls, as seen in the scRNA-seq data (Fig. 1I-K). Spatial plots (Fig. 4A, D and E) showed that increased immature oligodendrocytes and OPCs were specifically localized to the dorsal V-SVZ/corpus callosum where there was also a coincident decrease in mature oligodendrocytes. Quantification confirmed this conclusion; within the dorsal V-SVZ and corpus callosum, OPCs were increased approximately two-fold (79 +/- 9 to 159 +/- 54 SEM) and immature oligodendrocytes 3 to 4-fold (21 +/- 1 and 73 +/- 28 SEM) while mature oligodendrocytes were decreased by almost half (from 1370 +/- 152 to 713 +/- 137 SEM).

The second major difference involved microglia. UMAP visualization (Fig. 3F, G) identified two microglial clusters in the merged Xenium dataset. One cluster (20) included cells from both the control and remyelinating V-SVZ and corpus callosum. However, the other cluster (1) was almost completely comprised of cells from the remyelinating sections (92%). Cluster 20, was differentially-enriched for mRNAs defined for the homeostatic microglia such as *Ecscr* while cluster 1 was enriched for genes we had defined for the remyelination-enriched microglia such as *Cybb* (Fig. 4G; Suppl. Fig. 4D). Spatial plots (Fig. 4H, I; Suppl. Fig. 4D) showed that the large increase in transcriptionally-distinct microglia expressing genes like *Cybb* and *H2-Eb1* was largely limited to the dorsal V-SVZ and corpus callosum. Thus, the dorsal but not lateral V-SVZ NSC niche is altered during remyelination.

We confirmed these findings with a second type of multiplexed *in situ* hybridization-based single cell spatial transcriptomics, MERFISH^34–36^. We analyzed sections from the same and additional control and remyelinating brains, using a probeset targeting 300 genes (Suppl. Table 2). 119 of these targeted genes overlapped with the Xenium probeset, but the remaining 181 were distinct (see Suppl. Table 2). Nonetheless, in spite of the differing probesets, we obtained results similar to the Xenium analysis. Specifically, UMAP visualization and cell type-specific marker gene overlays (Fig. 4J; Suppl. Fig. 4E-G) identified the expected cell types in control and remyelinating conditions and spatial plots (Fig. 4K-M) showed that these were in the same locations as defined by Xenium. Moreover, the control dorsal and lateral NSC neighborhoods were distinct, with dorsal NSCs next to oligodendrocytes, and lateral NSCs next to neuroblasts and striatal neurons (Fig. 4L). Finally, the dorsal V-SVZ was specifically changed during remyelination, with a robust increase in dorsally-localized transcriptionally-distinct microglia (Fig. 4M; Suppl. Fig. 4H). This was coincident with an increase in OPCs and immature oligodendrocytes and a decrease in mature oligodendrocytes specifically in the dorsal V-SVZ and corpus callosum (Fig. 4K, L).

### Identification of niche-specific ligands predicted to act on NSCs in the control and remyelinating V-SVZ

These spatial findings describe differences in the V-SVZ NSC neighborhoods with regard to their cellular composition. We next asked how these cellular differences might influence the local NSC ligand environment. To do so, we first characterized receptors expressed by NSCs that could respond to local ligands. We used our scRNA-seq dataset and a previously-published ligand-receptor database^37^, defining a receptor mRNA as expressed if it was detectable in at least 5% of aNSCs or dNSCs. This analysis (Fig. 5A, Suppl. Table 3) identified 115 NSC receptor mRNAs, with 109 common to both control and remyelinating conditions. These common receptor mRNAs encoded previously-characterized NSC receptors such as TrkB/Ntrk2, PDGFRB, EDNRB, NOTCH1 and EGFR. Two receptor mRNAs were only detected in control NSCs (*Cxcr2* and *Vipr2*), and another five in remyelinating NSCs (*Prlr, Oprk1, Cxcr6, Cd74, Calcrl*). However, these condition-specific receptor mRNAs were expressed at very low relative levels, and differences might thus be attributable to small differences in dataset sensitivity.

**Figure 5.**
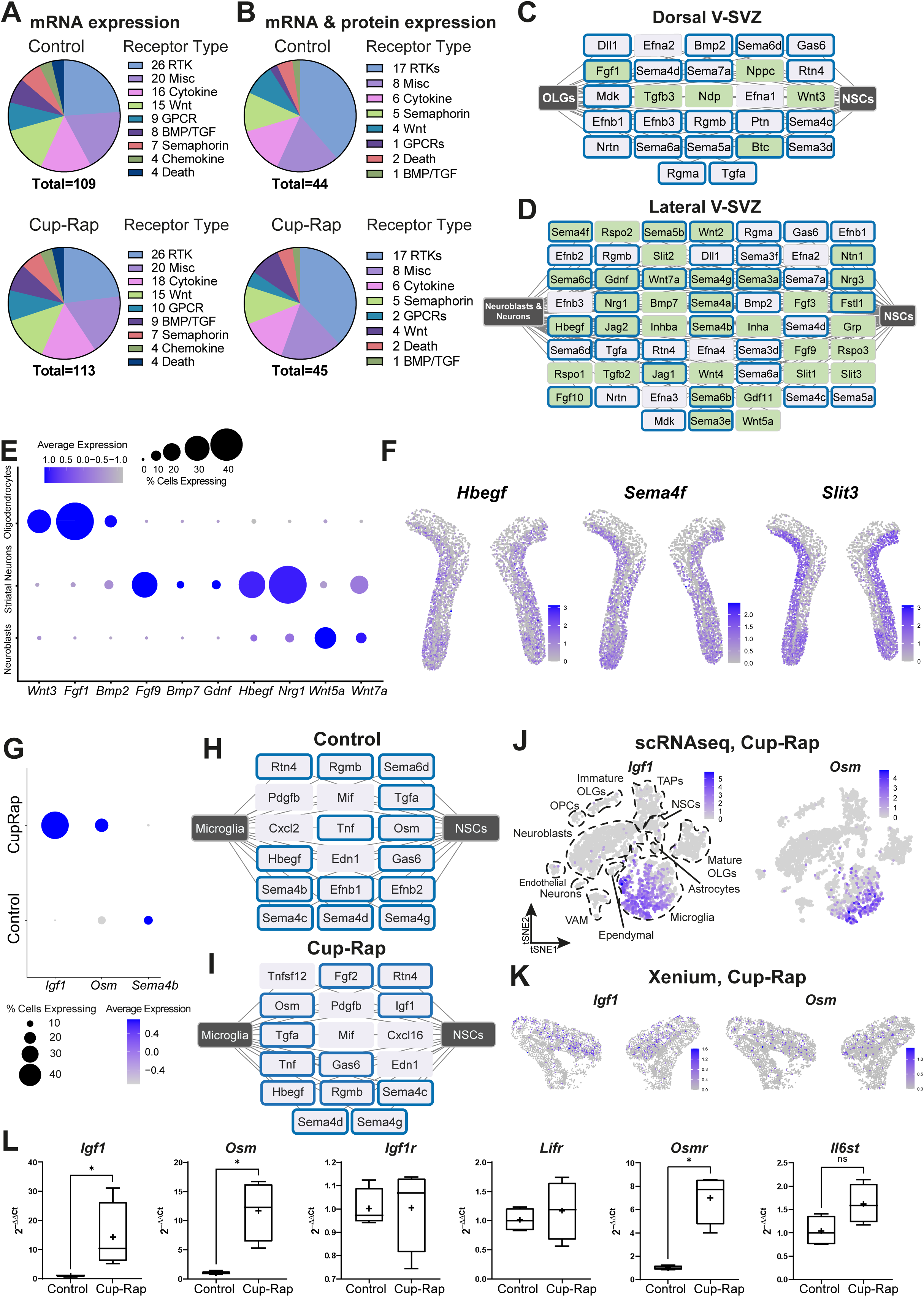
Analysis of the NSC ligand environment identifies niche-specific ligands predicted to act on NSCs in the control and remyelinating V-SVZ. Also see Suppl. Figure 5 and Suppl. Tables 3-6. (A) Pie chart classifying ligand-dependent receptors that were expressed at the mRNA level in NSCs from the control versus remyelinating V-SVZ, as defined from the merged CD1 vehicle and cuprizone-rapamycin-treated datasets (Fig. 1F, G) (also see Suppl. Table 3). **(B)** Pie chart classifying the 44 and 45 transcriptionally-identified control and remyelinating NSC receptors also detected using cell surface spectrometry of adult neurospheres (see Suppl. Table 4). **(C, D)** Models showing predicted interactions between control NSC receptors and ligands expressed by oligodendrocytes in the dorsal V-SVZ (C) or neuroblasts and proximal striatal neurons in the lateral V-SVZ (D). Each box represents a ligand and blue outlining indicates that the cell surface receptor was detected in the proteomic analysis. Green denotes ligands predicted to act on NSCs specifically in the dorsal or lateral V-SVZ niche, as relevant. **(E)** Dot plot of selected ligand mRNAs predicted to be highly enriched in the dorsal versus lateral V-SVZ showing relative expression in oligodendrocytes, neuroblasts and proximal striatal neurons, as determined from the scRNA-seq data (Fig. 1F and G). The size of the dot indicates the percentage of cells within a given cell type that express the mRNA, and the color indicates relative expression level, coded as per the adjacent key. **(F)** Spatial plots for three ligand mRNAs predicted to be highly enriched in the lateral V-SVZ in the model shown in (D). Relative cellular expression is shown in the V-SVZ and corpus callosum of a control section as analyzed using Xenium. Relative expression levels are coded as per the adjacent keys. **(G)** Dot plot showing expression of selected ligand mRNAs in microglia from the control versus cuprizone-rapamycin-treated scRNA-seq datasets. The size of the dot indicates the percentage of microglia detectably expressing the mRNA, and the color indicates relative expression level, coded as per the adjacent key. **(H, I)** Models showing predicted interactions between control (H) or remyelinating (I) NSC receptors and ligands expressed by microglia in the control (H) or remyelinating (I) V-SVZ. Each box shows a microglial ligand, and blue outlining indicates that the cell surface receptor was detected in the proteomics. **(J)** Gene expression overlays for *Igf1* and *Osm* on the merged CD1 mouse cuprizone-rapamycin-treated dataset (Fig. 1G), with expression levels coded as per the adjacent keys. **(K)** Spatial plots for *Igf1* and *Osm* mRNA expression in the dorsal V-SVZ, corpus callosum and top of the lateral V-SVZ on a representative remyelinating section, as determined from the Xenium-based analysis. Both lateral ventricles are from the same section. Relative expression levels are coded as per the adjacent keys. **(L)** Box plots showing the relative expression of mRNAs encoding IGF1, OSM and their receptors in three week recovery control or cuprizone-rapamycin treated V-SVZ tissue, as analyzed by qPCR. Relative expression was calculated using the ΔΔCt method with *Tbp* housekeeping gene. See *Methods* for primer details. Data are presented as boxplots with ‘Tukey’ whiskers, and crosses indicate sample means.

We asked which receptors were present as cell surface proteins, analyzing three independent replicates of cultured adult V-SVZ neurospheres to do so. These neurospheres are largely comprised of activated NSCs and TAPs, and can be cultured in sufficient numbers for proteomic analysis. Briefly, as previously described^38^, we performed periodate oxidation of cell-surface glycans, coupled the glycosylated proteins to a hydrazide resin^39,40^, digested them with trypsin, released the glycopeptides with PNGase F, and identified peptides by mass spectrometry. Of the NSC receptor mRNAs identified transcriptomically, 45 or approximately 40% were also detected by mass spectrometry (Fig. 5B; Suppl. Table 4). These included TrkB (Ntrk2), Notch1, LIFR, EGFR, CNTFR, IGF1R, EDNRB, IFNGR1, FGFR1 and GFRA1. The receptor repertoires were similar in the two conditions, with the exception of CALCRL, which was only detectably expressed in remyelinating NSCs.

We then asked how the control dorsal and lateral NSC niches might differ with regard to ligands for these receptors. We did not analyse cells common to both neighborhoods such as ependymal cells or astrocytes, since we found no evidence for spatial heterogeneity in these cell types, and thus assumed they would contribute similar ligands to both niches. Instead, we focussed on cell types specific to one or the other niche; oligodendrocytes for the dorsal V-SVZ and neuroblasts and striatal neurons for the lateral V-SVZ. To define ligands expressed by these cells, we analyzed the V-SVZ scRNA-seq dataset as we had for the NSC receptors. Of note, our dissection protocol ensured that any striatal neurons within these datasets must be very proximal to the V-SVZ. This analysis identified 39, 33 and 84 ligands expressed by oligodendrocytes, neuroblasts and proximal striatal neurons, respectively (Suppl. Table 5). We then modelled potential interactions between NSCs and oligodendrocytes for the dorsal V-SVZ, and neuroblasts plus proximal striatal neurons for the lateral V-SVZ. These analyses (Fig. 5C, D; Suppl. Table 6) predicted 27 dorsal neighborhood ligands and 59 lateral neighborhood ligands, with 21 ligands shared between the two environments. Notably, 6 ligands were specific to the dorsal niche (Fig. 5D, E), and two of these, *Fgf1*^41,42^ and *Wnt3a*^20^, have been reported to promote remyelination and oligodendrogenesis, respectively. By comparison, 38 ligands were specific to the lateral niche. Many of these were guidance receptors such as the Semaphorins and Slits, but others were well-known for enhancing adult neurogenesis, including *Fgf9, Bmp7, Gdnf, Hbegf, Neuregulin-1, Wnt5a* and *Wnt7a*^43–52^ (Fig. 5D, E). Probes for several of these ligands were included in the Xenium or MERFISH probesets and spatial plots for three of these, *Hbegf*, *Sema4f* and *Slit3* confirmed their localization in the lateral V-SVZ neighborhood (Fig. 5F). Thus, the dorsal versus lateral V-SVZ neighborhoods are differentially enriched for pro-glial versus pro-neurogenic ligands, consistent with cell genesis in these two domains.

We next asked how the dorsal NSC ligand environment was influenced by the large alteration in microglia in control versus remyelinating conditions. We again used the scRNA-seq datasets to identify microglial ligand mRNAs, considering those expressed in at least 5% of control and/or remyelinating microglia. This analysis identified 59 ligand mRNAs, including well-known microglial-associated ligands such as *Tnf, Il1a* and *Ccl3* (Suppl. Table 5). Notably, some ligands such as *Igf1* and *Osm* were more highly expressed during remyelination while others such as *Sema4b* were higher in uninjured controls (Fig. 5G).

We used these ligands to model interactions between microglia and NSCs in the control versus remyelinating dorsal V-SVZ (Figs. 5H, I; Suppl. Table 6). This analysis predicted 17 or 18 ligand-receptor interactions in the two conditions, with most involving receptors identified at both the transcriptomic and proteomic levels. Many predicted interactions were the same in both conditions, but two involving *Igf1* and *Fgf2* were predicted only in the remyelinating condition. Moreover, some interactions involved ligands that were significantly more highly expressed during remyelination, such as *Osm* (Fig. 5G-I).

### Two microglially-expressed ligands in the remyelinating dorsal NSC niche, IGF1 and OSM, promote NPC proliferation and oligodendrogenesis in culture

We asked whether any of the predicted microglially-expressed ligands had the potential to enhance oligodendrogenesis during remyelination. We focused on ligands that were specific to microglia and enriched in the remyelinating dorsal V-SVZ. Only two ligands met these criteria, IGF1 and OSM; both were specific to microglia (Fig. 5J), and both were enriched in the remyelinating dorsal V-SVZ niche (Fig. 5K). qPCR of isolated V-SVZ tissue (Fig. 5L) confirmed that *Igf1* and *Osm* mRNAs were increased during remyelination, in agreement with the scRNA-seq data (Fig. 5G). This analysis also confirmed expression of the relevant receptors in the V-SVZ, *Igfr1*, *Lifr, Osmr* and *Il6st/gp130*.

We asked if IGF1 or OSM could enhance NPC-dependent oligodendrogenesis using cultures of perinatal cortical precursors that appropriately generate postnatal V-SVZ progeny, oligodendrocytes and olfactory bulb interneurons^53^. Analysis of our previously-published scRNA-seq datasets^26^ confirmed that perinatal V-SVZ NPCs and OPCs detectably express *gp130, Lifr,* and *Igf1r* mRNAs (Suppl. Fig. 5A). To ensure we could measure changes caused by IGF1 or OSM precursors were cultured in 15 ng/ml rather than maximal 50 ng/ml FGF2. 15 ng/ml FGF2 supported cell survival as effectively as 50 ng/ml, but proliferation was reduced by about half (Fig. 6A).

**Figure 6.**
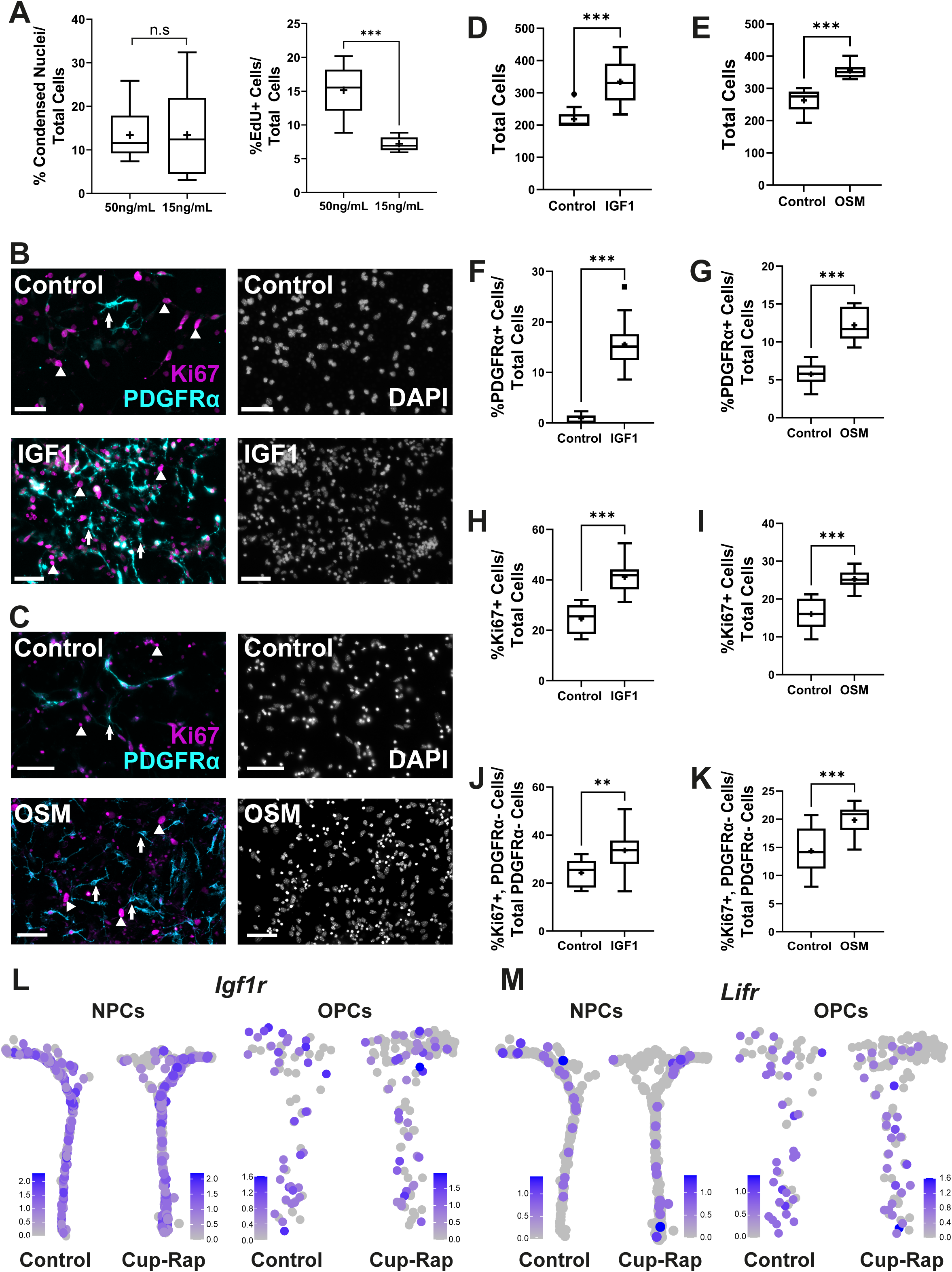
IGF1 and OSM promote NPC proliferation and genesis of OPCs in E15/16 cortical precursor cultures. Also see Suppl. Fig. 5. (A) E15/16 cortical precursor cells were plated in 15 or 50 ng/ml FGF2, 1 µM EdU was added after 3 days and cultures were analyzed after one further day by visualizing EdU and counterstaining with DAPI. Box plots show the percentage of cells with condensed, apoptotic nuclei (left panel) and the percentage of EdU-positive cells (right panel) in the cultures. N = 3 independent culture, 2 technical replicates per culture. ***p<0.001, ns = p>0.05. (B-K) E15/16 cortical precursor cells were plated in 15 ng/ml FGF2 with or without the addition of 100 ng/ml IGF1 (B, D, F, H, J) or OSM (C, E, G, I, K), and analyzed by immunostaining 4 days later. (B, C) Representative images of precursor cultures treated with IGF1 (B) or OSM (C) and immunostained for Ki67 (magenta) and PDGFRα (blue), and counterstained with DAPI (white). Arrows and arrowheads denote cells that are only positive for PDGFRα or Ki67, respectively. Scale bars = 50 µm. (D, E) Quantification of images similar to those in (B, C) for the average number of cells in four 800 µm^2^ randomly-selected quadrants treated with FGF2 alone or with FGF2 plus IGF1 (D) or OSM (E). For OSM, N = 3 independent cultures, 3 technical replicates per condition from each culture. For IGF1, N = 5 independent cultures, 3 technical replicates per condition from each culture. ***p<0.001. (F-K) Quantification of cortical precursor cultures treated with FGF2 alone (control) or with FGF2 plus IGF1 (F, H, J) or OSM (G, I, K) for the percentages of (F, G) OPCs expressing PDGFRα, (H, I) proliferating, Ki67-positive cells, or (J, K) proliferating neural precursors that were Ki67-positive and PDGFRα-negative, determined from images similar to those in (B, C). For OSM, N = 3 independent cultures, 3 technical replicates per condition from each culture. For IGF1, N = 5 independent cultures, 3 technical replicates per condition from each culture. **p<0.01, ***p<0.001. (L, M) Spatial plots for *Igf1r* (L) or *Lifr* (M) expression in NPCs or OPCs of the V-SVZ and corpus callosum, shown on representative control and cuprizone-rapamycin treated sections, as determined from the Xenium-based analysis. The NPCs and OPCs are grey dots and mRNA expression is overlaid in purple, coded as per the adjacent keys.

We then asked about IGF1 and OSM. We added 15 ng/ml FGF2 plus 100 ng/ml IGF1 or OSM upon plating, replenished the IGF1 or OSM at three days, and performed analysis one day later (4 days total). Quantification showed that IGF1 and OSM increased the total number of DAPI-positive cells within the cultures by approximately 1.5 and 1.4-fold, respectively (Fig. 6B-E). Immunostaining for PDGFRα and Ki67 (Fig. 6B, C) showed that in control cultures approximately 1 to 6% of total cells were PDGFRα-positive OPCs, and that this was significantly increased to almost 15% by either IGF1 or OSM (Fig. 6F, G). Both ligands also significantly increased proliferating Ki67-positive cells (Fig. 6B, C, H and I), consistent with the increased total cell numbers. The majority of proliferating cells were Ki67-positive, PDGFRα- negative NPCs and these were significantly increased by IGF1 and OSM (Fig. 6J, K). Proliferating PDGFRα-positive OPCs were also observed, although numbers were low, particularly in the control FGF2 only conditions (mean 3.4 +/- 0.6 SEM proliferating OPCs across controls from both conditions). Thus, both IGF1 and OSM promote the proliferation of V-SVZ NPCs and enhance the genesis of OPCs within these cultures.

### IGF1 and OSM cause expansion of NPCs and OPCs in vivo

We asked if IGF1 or OSM would also stimulate expansion of NPCs or OPCs *in vivo*, using minipumps to infuse these ligands directly into the lateral ventricles of adult mice for one week (Suppl. Fig. 5B). Spatial plots of the Xenium data confirmed that *Igf1r* and the OSM receptor *Lifr* were expressed in NPCs and OPCs around the lateral ventricles of control mice (Fig. 6L, M). A second OSM receptor, *Osmr* was detectable only in a small number of V-SVZ NPCs (data not shown), reflecting its much lower expression in the scRNA-seq datasets (for example, see Suppl. Fig. 5A).

We analyzed coronal forebrain sections through the V-SVZ and corpus callosum immediately after growth factor infusion by immunostaining for Sox2 to detect NPCs, PDGFRα to detect OPCs, and Ki67 to detect proliferating cells (Fig. 7A, G). Quantification of the dorsal wall of the V-SVZ and the corpus callosum showed that IGF1 significantly increased the numbers of Sox2-positive, PDGFRα-negative NPCs in this region by about 1.5-fold (Fig. 7A,B). The number of PDGFRα-positive OPCs was also increased by more than 2-fold (Fig. 7A, C). These increases were at least partly due to enhanced proliferation, since the total number of Ki67-positive cells was significantly increased (Fig. 7A, D), as were Ki67-positive, Sox2-positive NPCs and Ki67-positive, PDGFRα-positive OPCs (Fig. 7E, F).

**Figure 7.**
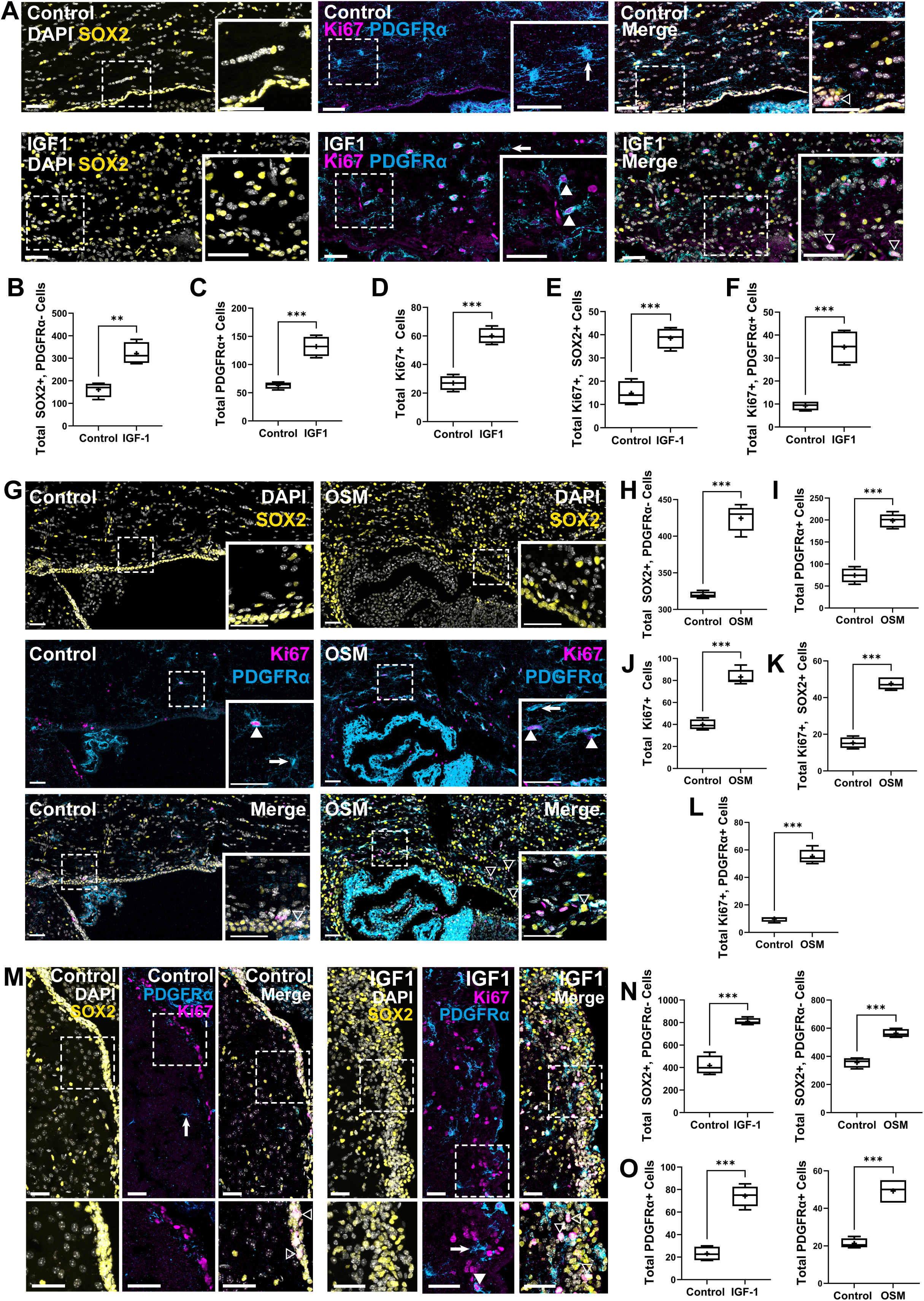
Infusion of IGF1 or OSM into the lateral ventricles promotes adult NPC proliferation and oligodendrogenesis in vivo. Also see Suppl. Figure 5. IGF1 (A-F, M-O), OSM (G-L, N, O) or 0.1%BSA/PBS (Control) were infused into the lateral ventricles of control CD1 mice for one week using minipumps, and coronal forebrain sections through the V-SVZ and corpus callosum were analyzed immediately after by immunostaining (see Suppl. Fig. 5B). **(A, G)** Representative high magnification confocal images of the dorsal V-SVZ and corpus callosum from mice infused with vehicle (Control), IGF1 (A) or OSM (G), immunostained for Sox2 (yellow), PDGFRα (blue), and Ki67 (magenta). Also shown is DAPI counterstaining to highlight nuclei (white). The boxed regions are shown at higher magnification in the insets. Arrows, filled arrowheads and empty arrowheads denote cells positive for only PDGFRα, PDGFRα and Ki67, and Sox2 and Ki67, respectively. Scale bars = 100 µm (tiled image) and 50 µm (insets) **(B, H)** Quantification of images similar to those in (A, G) for total Sox2-positive, PDGFRα-negative NPCs, counting cells along the entire medial-lateral axis of the V-SVZ dorsal wall and adjacent corpus callosum in each of two sections per animal. N = 8 animals per condition, **p<0.01, ***p<0.001. **(C, I)** Quantification of images similar to those in (A, G) for total PDGFRα-positive OPCs, counting cells along the entire medial-lateral axis of the V-SVZ dorsal wall and adjacent corpus callosum in each of two sections per animal. N = 8 animals per condition, ***p<0.001. **(D, J)** Quantification of images similar to those in (A, G) for total Ki67-positive proliferating cells, counting cells along the entire medial-lateral axis of the V-SVZ dorsal wall and adjacent corpus callosum in each of two sections per animal. N = 8 animals per condition, ***p<0.001. (**E, K)** Quantification of images similar to those in (A, G) for total Ki67-positive, Sox2-positive proliferating NPCs, counting cells along the entire medial-lateral axis of the V-SVZ dorsal wall and adjacent corpus callosum in each of two sections per animal. N = 8 animals per condition, ***p<0.001. **(F, L)** Quantification of images similar to those in (A, G) for total Ki67-positive, PDGFRα-positive proliferating OPCs, counting cells along the entire medial-lateral axis of the V-SVZ dorsal wall and adjacent corpus callosum in each of two sections per animal. N = 8 animals per condition, ***p<0.001. **(M)** Representative high magnification confocal images of the lateral V-SVZ of mice infused with IGF1 or vehicle (Control) and immunostained for Sox2 (yellow), PDGFRα (blue), and Ki67 (magenta). Also shown is DAPI counterstaining to highlight nuclei (white). The boxed regions are shown at higher magnification in the bottom panels. Arrows, filled arrowheads and empty arrowheads denote cells positive for only PDGFRα, PDGFRα and Ki67, and Sox2 and Ki67, respectively. Scale bars = 100 µm (tiled image) and 50 µm (insets) **(N, O)** Quantification of images similar to those in (M and Suppl. Fig. 5B) for total Sox2-positive, PDGFRα-negative NPCs (N) and total PDGFRα-positive OPCs (O), counting cells along the entire dorsal-ventral axis of the V-SVZ lateral wall in each of two sections per animal. Cells were included if they were within 150 µm of the ventricle wall. N= 8 per animals per condition, ***p<0.001.

Similar results were obtained with OSM infusion (Fig. 7G). The numbers of Sox2-positive, PDGFRα-negative NPCs and PDGFRα-positive OPCs in the dorsal wall/corpus callosum were significantly increased by about 1.3 and 2.7-fold, respectively (Fig. 7G-I). Moreover, the numbers of total Ki67-positive cells, Ki67-positive, Sox2-positive NPCs and Ki67-positive, PDGFRα-positive OPCs were all significantly increased (Fig. 7J-L).

These data indicate that locally-increased IGF1 and OSM can enhance genesis of OPCs in the dorsal wall/corpus callosum, a region where NPCs normally generate glial cells. Since lateral wall NPCs also express receptors for these two ligands even though they normally generate neuroblasts (Fig. 6L, M), we asked if oligodendrogenesis was also altered in this region. Quantification of immunostained sections revealed that IGF1 and OSM both significantly increased Sox2-positive, PDGFRα-negative NPCs, and PDGFRα-positive OPCs in the lateral V-SVZ (Fig. 7M-O; Suppl. Fig. 5C). Thus, these two growth factors can enhance oligodendrogenesis even in a V-SVZ region where NSCs normally make neurons.

## DISCUSSION

The idea that myelin repair can be promoted by recruitment of endogenous neural precursors is an exciting one. However, to realize this therapeutic vision, we need to understand both the precursors themselves, and their local environments under homeostatic and pathological conditions. Here, we have used lineage tracing, scRNA-seq and single cell spatial transcriptomics to provide a molecular and cellular overview of adult forebrain NSCs and their niches during homeostasis and remyelination. We show that in the uninjured brain the dorsal and lateral V-SVZ comprise very different cellular niches for NSCs, consistent with the differential genesis of glia versus olfactory bulb interneurons in these two regions. We also show that during remyelination, only the dorsal niche is altered, with a robust local increase in transcriptionally-distinct microglia, coincident with increased NPC-dependent oligodendrogenesis. We define ligands within these different NSC neighborhoods that could act on NSC receptors, and identify IGF1 and OSM as ligands that are locally increased in microglia in the dorsal V-SVZ during remyelination. These ligands cause proliferation of NPCs and increased oligodendrogenesis both in culture and when infused into the lateral ventricles, even in the lateral V-SVZ niche where NSCs normally make interneurons. These data support a model where gliogenesis versus neurogenesis is determined by the local NSC neighborhood and where injury-induced niche alterations promote NSC activation, local oligodendrogenesis, and potentially myelin repair.

Previous studies have shown that the forebrain NSC niche is a mosaic, with NSCs in the dorsal V-SVZ preferentially generating glial cells, and those in the lateral V-SVZ olfactory bulb interneurons (reviewed in ^11–13^). One potential explanation for this finding is that NSCs within these domains have distinct differentiation potentials conferred by their developmental origin, since NSCs in the dorsal wall are cortically-derived, while most lateral wall NSCs are GE-derived^33,54–58^. In this regard, we previously showed that cortical and GE-derived NSCs do retain a transcriptional memory of their developmental origins, but are nonetheless highly similar transcriptionally^26^. Moreover, both cortically and GE-derived NSCs generate olfactory bulb interneurons and glial cells to some degree, arguing that developmental origin alone does not dictate genesis of neurons versus glia.

What then explains the mosaic nature of V-SVZ cell genesis? We propose that it is the NSC niche that differs. We show that, as predicted, NSCs in both the dorsal and lateral V-SVZ are closely-associated with ependymal cells, but our spatial transcriptomic data also highlight differences between these two niches; dorsal NSCs are closely associated with corpus callosum oligodendrocytes while lateral NSCs are closely associated with their neuroblast progeny and neighboring striatal neurons. Since oligodendroglia, striatal neurons and neuroblasts all express distinct ligands that can act on the NSC receptors we have defined here, then this implies that NSCs in these two neighborhoods are exposed to very different growth factor environments. Intriguingly, many of the lateral niche growth factors we have defined, including *Fgf9, Bmp7, Gdnf, Hbegf, Neuregulin-1, Wnt5a* and *Wnt7a* are known to directly promote neurogenesis^43–52^, while several of the dorsal niche ligands *Wnt3a* and *Fgf1* can promote oligodendrogenesis and remyelination, respectively^20,41,42^. These distinct ligand environments thus provide one mechanism for differential cell genesis in these two regions.

The mosaic nature of the V-SVZ niche environment is even more apparent following white matter injury. Together, our lineage tracing, scRNA-seq and spatial transcriptomic data identify increases in NPC-dependent oligodendrogenesis and microglia that are largely limited to the dorsal V-SVZ and corpus callosum. These locally increased microglia express two ligands, OSM and IGF1 that we show can act directly on NPCs to enhance oligodendrogenesis in culture and *in vivo*. Notably, NSCs within the remyelinating dorsal niche do not change their transcriptional profiles, but only their activation states, supporting the idea that it is the environment and not the stem cells that are fundamentally changing. We also show that IGF1 and OSM intraventricular infusion promote oligodendrogenesis in the lateral V-SVZ, an area that normally makes predominantly olfactory bulb neuroblasts, further supporting the idea that the niche environment plays a key role in determining cell genesis.

The distinct transcriptional state we document here for remyelinating dorsal microglia is similar to states previously described in other neural pathological situations, including experimentally-induced demyelination^59–61^. While immune activation is normally considered to be a negative event in disorders like multiple sclerosis^62,63^, in other types of injury, tissue-resident macrophages or microglia are thought to promote tissue repair (reviewed in ^64^), much as we are suggesting here. It is likely that in situations where repair is possible (such as cuprizone-rapamycin-induced remyelination), microglia play a predominantly positive role, but that when repair responses go awry, the more negative inflammatory aspects of these cells come to the fore.

One of the local remyelination-specific microglial ligands we identify here, IGF1, has previously been implicated in myelination^65–69^. Of particular relevance is a study^70^ identifying developing *Igf1*-expressing Cd11c-positive microglia and showing that conditional loss of *Igf1* in these cells negatively impacts developmental myelination. How then does IGF1 promote myelination since it seems to be important both during development and following injury? Previous studies have focused on IGF1’s effects on OPC survival and/or on oligodendrocytes^65–69^. Our data do not preclude these mechanisms, but instead show that IGF1 also promotes NPC proliferation and directly enhances oligodendrogenesis. Moreover, our data indicate that these effects are likely mediated, at least in part, by a local increase in microglial IGF1 secretion in the dorsal V-SVZ neighborhood.

The second microglial ligand we identify, OSM, is also specifically expressed by microglia, is robustly and locally increased during remyelination and promotes both NPC proliferation and oligodendrogenesis. Previous work has implicated OSM in modulating the immune response and astrocytes during remyelination^71,72^, but our data argue that it also directly regulates NPCs, likely by activating one of two OSM receptors, LIFR or OSMR. Intriguingly, LIFR is also an obligate receptor for the related ligands LIF, CNTF and cardiotrophin-1 and all of these enhance the genesis of glial cells from embryonic cortical precursors^73–75^. Moreover, both CNTF and LIFR downstream signaling regulate adult demyelination, an effect thought to be mediated in part via oligodendrocytes^76,77^. We therefore speculate that in homeostatic conditions LIFR signaling functions to regulate dorsal NSC-dependent gliogenesis, and that following white matter injury microglial OSM functions to hyperactivate this pathway and promote local remyelination.

One other key question is whether endogenous recruitment of NSCs for white matter repair represents a realistic strategy. While there is as of yet no definitive answer to this question, increasing evidence supports the idea. As one example, we previously identified metformin as a drug that enhances the differentiation of mammalian (including human) NSCs^1^, and various studies have since shown that metformin promotes brain and myelin repair in animal models^2,7,78,79^. Notably, a pilot clinical trial using metformin on children with an acquired brain injury due to radiation treatment of their brain tumors showed that metformin treatment was associated with several measures of cognitive recovery and white matter improvement^2^. Thus, if we can better-define the environmental cues that normally promote oligodendrogenesis, then perhaps this will ultimately provide us with more effective ways to promote endogenous brain repair.

## ACKNOWLEDGEMENTS

This work was funded by grants from the CIHR (to D.R.K. and F.D.M.), the CFREF “Medicine by Design” (to D.R.K. and F.D.M.), and the Canadian Stem Cell Network (to D.R.K. and F.D.M.). A.W. was funded by an M.S. Society of Canada fellowship and D.J. by CIHR and Hospital for Sick Children Restracomp studentships. We thank Rebecca Parsons, Nareh Tahmasian, Neenah Williams, Michelle Wojciechowski, and Jasmine Yang for help with some of the experiments.

## AUTHOR CONTRIBUTIONS

A.W. and D.J. conceptualized and analyzed experiments and co-wrote the paper. D.J. established the cuprizone-rapamycin model, performed the scRNA-seq and lineage tracing experiments and analyzed these data. A.W. performed the Xenium and MERFISH experiments and analyses, performed the functional experiments with IGF1 and OSM, analyzed scRNA-seq data and performed the qPCR analysis. Y.L. performed the OSM and IGF1 *in vivo* infusions. M.A.L. worked with A.W. to perform the culture experiments. S.Y. performed and analyzed the neurosphere mass spectrometry analysis. P.W.F. conceptualized the infusion experiments and co-wrote the paper. F.D.M. and D.R.K. conceptualized experiments, analyzed data, and co-wrote the paper.

## DATA AVAILABILITY

scRNA-seq, Xenium, and MERFISH datasets will be deposited in the GEO database and made available upon acceptance of this paper.

## EXPERIMENTAL METHODS

### Mice

All animal protocols were approved by the Animal Care Committees of either the Hospital for Sick Children or the University of British Columbia, as relevant, and were in accordance with the Canadian Council of Animal Care policies. Animals had free access to rodent chow and water and were housed in a temperature and humidity-controlled environment on a 12-hour light-dark cycle. Additional supportive care was provided for animals receiving cuprizone-rapamycin treatment, including hydration and heat support. All animals were regularly monitored, and mice displaying distress or health issues were provided with supportive measures or euthanized if a humane endpoint was reached. No animals included in our experiments displayed obvious motor or behavioural phenotypes and none were immune compromised. No animals included in our experiments lost more than 25% of their starting body weight.

Wild-type CD1, *NestinCreERT2* (C57BL/6-Tg(Nes-cre/ERT2))^80^, and *R26R^Eyfp/Eyfp^* (B6.129X1-Gt(ROSA)26Sortm1(EYFP)Cos/J)^81^ mice were all obtained from Jackson Laboratories. For lineage tracing experiments, *NestinCreERT2* heterozygous mice were crossed with *R26R^Eyfp/Eyfp^* mice, and the resultant *NestinCreERT2: R26R^Eyfp/Eyfp^* (denoted as *NestinCreErt2-Eyfp*) mice were analyzed. All mice were bred and genotyped as recommended by Jackson Laboratories.

For single cell RNA sequencing (scRNA-seq) studies using wild-type CD1 mice and subsequent RT-qPCR validation experiments, only females were used. For scRNA-seq and *in vivo* lineage tracing experiments involving *NestinCreErt2-Eyfp* mice, animals of either sex were used, and no sex dependent effects were observed. In all cases, mice were randomly allocated to experimental groups. For *in vitro* cortical precursor cell cultures, embryos were collected at embryonic day 15-16 (E15-16) from adult CD1 mothers. Embryo sex was undetermined prior to culture. For *in vivo* ligand infusion experiments, equal samples of CD1 mice of each sex were used and no sex-dependent effects were observed.

All *in vivo* mouse experiments used adult mice. Animals involved in demyelination experiments were aged 5-6 weeks (postnatal day 40-45 (P40-45)) at the beginning of cuprizone-rapamycin (or vehicle-control) treatment. Animals included in morphological lineage tracing studies were aged 8 weeks (P56). Animals were between 12-14-weeks-old (P84-98) for minipump ligand infusion experiments.

### Tamoxifen administration

For morphological lineage tracing experiments, adult 8 week-old *NestinCreErt2-Eyfp* animals were injected intraperitoneally with 3 mg of tamoxifen (Sigma) dissolved in a sunflower oil/ethanol mixture (9:1) to 30 mg/mL, once daily for five consecutive days. Mice were analyzed either 8 weeks later (Figure 1A, Suppl. Figure 1B) or immediately (Suppl. Figure 1A). For *NestinCreErt2-Eyfp* morphological lineage tracing experiments involving demyelination (Figure 1B-E), recombination was induced with tamoxifen during the first week of recovery from six weeks of cuprizone-rapamycin or vehicle-control treatment. Tamoxifen was prepared as above and administered intraperitoneally once daily for five consecutive days. Analysis of mice was performed 3 weeks after the cessation of cuprizone-rapamycin or control-vehicle treatment. For lineage traced scRNA-seq experiments, tamoxifen was administered once daily for five days prior to the start of the cuprizone-rapamycin or vehicle treatment protocols.

### Cuprizone-rapamycin treatment

To induce demyelination in adult mice, we used a cuprizone and rapamycin model as described in Sachs et al. 2014^24^ (see also ^28^). P40-45 CD1 or *NestinCreErt2-Eyfp* mice were fed *ad libitum* powdered chow containing 0.3% cuprizone (biscyclohexanone-oxaldihydrazone; C9012, Sigma) for six weeks. Intraperitoneal injections of rapamycin (10 mg/kg rapamycin, R-5000, LC Laboratories; dissolved in 5% ethanol, 5% polyethylene glycol 400 (PEG400), Sigma; 5% Tween 80, Sigma; in PBS) were also administered for five consecutive days per week for six weeks to inhibit spontaneous, endogenous remyelination from resident OPCs and NSCs. Age and sex matched controls were fed standard powdered mouse chow and received vehicle injections (5% PEG400; 5% Tween 80; and 5% ethanol, reconstituted in PBS) for the six-week period. Upon cessation of the cuprizone-rapamycin (or vehicle-control) treatment period, mice were collected either immediately with no recovery or after three weeks of recovery (approximated as the mid-point of remyelination in the cuprizone-rapamycin model^24^).

### Mini-pump implantation and ligand infusion

CD1 mice, aged 12-14 weeks (P84-98), were pre-treated with atropine sulfate (0.1 mg/kg, intraperitoneal), anesthetized with chloral hydrate (400 mg/kg, intraperitoneal), and administered the analgesic meloxicam (4 mg/kg, subcutaneous). Following this, the mice were placed in a stereotaxic frame, and lidocaine was topically applied around the incision site. Osmotic Minipumps (Model 1007D) and Brain Infusion Kits (0008851, Alzet Osmotic Pumps) were implanted at the following coordinates relative to bregma: AP 0 mm, ML +0.8 mm, DV -2.5 mm. IGF-I (RND Systems, 791-MG-050) or OSM (RND Systems, 495-MO-025) were diluted in PBS. Minipumps were filled with either IGF (34 μg/mL), OSM (17 μg/mL) or vehicle (0.1%BSA/PBS) and inserted subcutaneously for 7 days. Consequently, mice received either 408 ng of IGF or 204 ng of OSM per day for 7 days, with a delivery rate of 0.5 μL per hour. Upon infusion end point, mice were euthanized and cardiac perfusions with ice-cold PBS, followed by 4% paraformaldehyde (PFA) performed.

Brains were sucrose protected and flash frozen in Optimal Cutting Temperature (OCT) embedding matrix (TissueTek) for subsequent cryosectioning and immunohistochemical analyses. Eight mice (4 female) were assigned to each treatment or associated control group.

## METHOD DETAILS

### Single cell isolation and 10X Genomics sequencing

For each scRNA-seq run, tissues were collected from four to five non-littermate mice. Single cells were isolated from adult CD1 or *NestinCreErt2-Eyfp* V-SVZ tissue from either the demyelinated injury or control group as previously described^26^. Briefly, mice were euthanized and dorsal and lateral wall V-SVZ tissues were dissected in ice cold Hanks buffered salt solution (HBSS). Tissues were finely chopped and incubated in Hibernate A minus calcium medium (Brain Bits) supplemented with 2% B27, 20 units/mL papain (Worthington), and 0.005% DNaseI (Worthington) for 30 minutes at 37°C in the dark. This enzymatic incubation solution also contained DRAQ5, a membrane permeable dye to stain for live cells (at 5 μM or 3000-fold dilution, Thermo Fisher). After enzymatic treatment, the tissue suspension was centrifuged at 300 x g for 5 minutes and resuspended in HBSS/0.025% BSA. A cell suspension was produced via manual trituration using blunt needles of sequentially decreasing diameter (McMaster-Carr). The resultant cell suspension was filtered through a 40 μm strainer and labeled with 1 μg/mL propidium iodide (PI, Abcam). Viable cells were collected using flow cytometry sorting for propidium iodide-negative and DRAQ5-positive cells. For all experiments, single cell capture and cDNA library preparation were conducted at the Princess Margaret Genomics Centre (Toronto, ON) using the 10X Genomics Chromium system protocol.

### scRNA-seq data analysis

cDNA libraries were sequenced on Illumina HiSeq2500 at the Princess Margaret Genome Centre (Toronto, ON) with an average read depth of 65,000 reads per cell. FASTQ reads were then processed using CellRanger V3 software with recommended settings by the manufacturer (10X Genomics).

scRNA-seq data were analyzed according to our previously described computational pipeline^25–28,82^. Briefly, cells were excluded based on low UMI counts, predicted doublets, predicted contaminant red blood cells, and high mitochondrial DNA content. Genes which were detected in less than three cells were also removed. To correct for read depth and library size variation, cell transcriptomes were normalized using an iterative deconvolution normalization method (described in ^83^). The cell cycle phase was predicted for each cell using the Cyclone approach^30^. Normalized matrices were imported into Seurat and principal component analysis (PCA) performed using the top 2000 highly variable genes. Two-dimensional t-SNE projections were generated using the top principal components (RunTSNE, Seurat). The same principal components were also used to execute SNN-Cliq-inspired clustering (FindClusters Seurat) with iteratively increasing resolution until the number of differentially expressed genes (FindMarkers Seurat, where p < 0.01 family wise error rate (FWER), Holm method) between the most similar clusters reached around 30 genes. All individual or merged datasets were analyzed with the most conservative resolution selected based on clustering that best aligned with the expression of well-defined markers for each cell type. The selected resolution and number of clusters for each dataset can be found in its associated figure legend.

We used gene expression overlays on t-SNE plots (FeaturePlots, Seurat) to annotate cell types and state based on the following well-defined marker genes, including: For all NSCs, *Rcn3, Nes, Veph1, Notum, Tspan18*; for activated NSCs, *Egfr, Ascl1*; for dormant NSCs, *Meg3, Sparc, Fbxo2, Id3*; for TAPs, *Mki67, Rrm2, Hells, Gsx2*; for astrocytes, *Aqp4, Agt, Hsd11b1, Lcat*; for all oligolineage cells: *Olig1, Sox10* ; for immature oligodendrocytes, *Enpp4*; for OPCs, *Pdgfra, Cspg4*; for mature oligodendrocytes, *Mog, Mag, Opalin;* for all neuroblasts, *Dlx1, Dlx2, Sp8, Sp9, Gad1, Gad2*; for all microglia, *Aif1, Trem2, Cx3cr1, Tmem119*; for other immune cells including T cells and neutrophils, *Cd52*, *Cd69*; for striatal neurons, *Calb1, Gad1, Gad2, Bcl11b*; for endothelial cells, *Pecam1, Esam, Plvap*; for pericytes, *Carmn, Cspg4, Ano1*; for VAMCs, *Pdgfrb, Myh11, and Mylk*; and for ependymal cells, *Foxj1, Pvalb*.

Following this initial analysis of full individual datasets, merges and subsets were generated. To this end, barcodes of cells of interest were used to extract gene expression information from their associated filtered gene expression matrices. For merged datasets, the extracted gene expression information from each dataset was combined. Subsets and/or merges of cells were then run through the previously described pipeline once more. For all scRNA-seq analyses, Seurat versions were minimum v3.2.3 and maximum v.5.0.1.

### Batch correction of scRNA-seq data

To reduce batch effects from CD1 and *NestinCreErt2-Eyfp* datasets, batch correction (Suppl. Figure 2D) using the R package Harmony (v1.2.0)^29^ was performed. Using the RunHarmony function, one iteration of batch correction on the merged Seurat object containing all neural cells from CD1 and *NestinCreErt2-Eyfp* was performed. Following this correction, new t-SNE embeddings were generated and the data were re-clustered at various resolutions using the FindNeighbors and FindClusters functions, as detailed in the previous section.

### Differential gene expression and creation of gene signatures

To investigate transcriptional differences and create gene signatures for our observed homeostatic and remyelination-enriched microglia, differential gene expression analysis was performed. This analysis was performed using the Seurat FindMarkers function on the merged CD1 control and cuprizone microglia subset (Figure 3A) to compare clusters 0, 1, 3, 6, 10, 11 with clusters 2, 4, 5, 7, 8, 9, 12. Differentially enriched gene lists were refined by including only genes with a Bonferroni adjusted p-value < 0.05 (Adj. p-value) and ≥ 1.0 log2 fold-change in average expression which were expressed in >10% of cells in either the homeostatic or remyelination-specific group (dependent on comparison directionality).

Gene signatures used for scRNA-seq and subsequent *in situ* based spatial transcriptomics analyses were refined using t-SNE gene expression overlays to assess expression level and specificity. The homeostatic gene signature included 39 genes which were: *Adgrg1, Adrb1, Arhgef40, Asb2, Bank1, Ccnd1, Ccr5, Ccr6, Chst7, Crybb1, Csmd3, Dok3, Ecscr, Fam110a, Fchsd2, Gimap6, Gpr34, Il7r, Jam2, Kcnk12, Lrba, Med12l, Myo1b, Nav3, Numb, P2ry10b, P2ry12, P2ry13, Plxna4, Sall1, Selplg, Serpinf1, Slc2a5, Slc40a1, Slco4a1, Srgap2, Tmem100, Tmem119, Tspan18*. The remyelination enriched signature included 64 genes which were: *Atp6v0d2, Ass1, Klrb1a, Ms4a7, Gpnmb, Gpx3, H2-Eb1, Wfdc17, Msr1, Dcstamp, Acp5, Klrb1b, H2-Aa, Lgals3, Fn1, Il3ra, Bhlhe40, Apoc4, Rab7b, Igf1, AB124611, Bambi, Colec12, Vim, Adam33, Cybb, Gas7, Anxa2, Alcam, Itga4, Flt1, Cxcl14, Cst7, H2-Ab1, Atf5, Fabp3, Ahnak2, Atp1a3, Iqgap1, Gm1673, Mdfic, Rai14, Cd22, Cyb5r1, Ifi202b, Itgax, Crip1, Ifitm3, Cox6a2, Capg, Fabp5, Chst2, Ctse, Lpl, Cdkn2a, Fxyd5, Spp1, 1700003F12Rik, Anxa5, Lgals1, Apoc1, Cd74, Axl, Lyz2*.

### scRNA-seq and proteomic-based predictive ligand-receptor communication models

To investigate the V-SVZ niche microenvironment and interrogate potentially important ligand-receptor interactions with NSCs, predictive models of ligand and receptor communication were generated from scRNA-seq datasets using methods previously described^37,38^. Communication models of the dorsal and lateral V-SVZ were generated by first extracting ligand mRNAs expressed by cell types enriched in either dorsal or lateral regions, as informed by our single cell spatial transcriptomic data. For the dorsal V-SVZ model, we extracted all ligand mRNAs expressed by oligodendrocytes, which were enriched in the corpus callosum adjacent to the dorsal V-SVZ (Figure 4A, E and F). For the lateral V-SVZ model, we extracted all ligand mRNAs expressed by neuroblasts and striatal neurons which were predominant in the lateral V-SVZ and surrounding region (Figure 4A, B and F). Given the significant increase and transcriptional distinctions we observed in microglia between control and remyelinating conditions (Figure 1H, Figure 3A-E), we also produced predictive models of microglial ligand-NSC receptor interactions using datasets from vehicle-control and cuprizone-rapamycin treated mice.

For regional models of the V-SVZ at homeostasis, all ligand data were extracted from the CD1 control datasets, except for striatal neuron ligands which were not present in these datasets (Suppl. Table 5). Striatal neuron ligand mRNA expression data were instead extracted from the *NestinCreErt2-Eyfp* control datasets. For microglia, where we predicted interactions in both control and remyelinating conditions, data were extracted from CD1 vehicle-control and cuprizone-rapamycin datasets and matched with receptor data from NSCs derived from the same condition (i.e. control microglia ligands and control NSC receptors). Receptor mRNAs expressed by NSCs were extracted from CD1 vehicle-control and, in the case of the microglia-NSC model, cuprizone-rapamycin datasets (Suppl. Table 3). All ligands and receptors were extracted based on our previously published curated ligand and associated receptor database^37^.

Following initial ligand and receptor list extraction, percentages of cells expressing each ligand mRNA were calculated for each cell type. Ligands were defined as expressed if they were detected in at least 5% of cells. For dorsal and lateral V-SVZ models, NSC receptors were defined as expressed if they were expressed by at least 5% of activated and/or dormant NSCs in the CD1 control dataset. For microglia ligand-NSC receptor models, NSC receptors were defined as expressed if their mRNAs were expressed in at least 5% of activated or dormant NSCs in either the vehicle-control or cuprizone-rapamycin dataset. Ligand and receptor interactions were included in our communication models if both the ligand and associated receptor(s) met these criteria (Suppl. Tables 3, 5, and 6).

Cytoscape (v3.9.1) was used to visualize these predictive ligand-receptor communication models, where ligands included in the model are presented in a central panel of nodes and edges connecting them to their source and target cell type. Cell surface mass spectrometry analysis of V-SVZ neurospheres was incorporated into the models (see *Preparation of cultured V-SVZ neurospheres for cell surface mass spectrometry* and Suppl. Table 4 for further detail).

### Preparation of cultured V-SVZ neurospheres for cell surface mass spectrometry

Periventricular tissue was dissected and cultured from N = 3 male 6–8-week-old CD-1 mice as previously described^84^. Briefly, single-cell suspensions of dissected tissue were prepared, plated in one well of a 6-well plate per brain and allowed to grow for one week to form primary spheres. Subsequently, all primary spheres were passaged into one T175 flask (Sarstedt) by mechanical dissociation with a P1000 pipette and straining through a 70 µm cell strainer. One week later following secondary sphere growth, spheres were transferred to a 50 ml tube and spun down at 110 x g. Spheres were then resuspended in coupling buffer (1x PBS pH 6.5, 0.1% FBS) treated with 5 mM NaIO4 in coupling buffer for 30 min at room temperature in the dark and then processed as previously described^38^, with the exception that data presented in Suppl. Table 4 are spectral counts for each identified protein (identified as protein false discovery rate < 0.01) in each of the three replicates.

### RNA isolation, cDNA synthesis and Reverse Transcription-Quantitative PCR (RT-qPCR)

For RT-qPCR validation of ligand and receptors identified by communication modeling, dorsal and lateral V-SVZ tissue was collected from adult CD1 mice which had been subject to either vehicle-control or cuprizone-rapamycin treatment. Tissues were dissected immediately following euthanization in ice-cold HBSS and stored in RNAlater (Sigma, R0901) for up to 2 weeks at -70°C. V-SVZ issues were placed in lysis buffer (Qiagen, 74104) and mechanically homogenized in individual 1 ml tubes containing RNase free 2.8 mm ceramic beads (Qiagen, 13114-50). RNA isolation was performed using RNeasy mini-kits according to the manufacturer’s protocol (Qiagen, 74104). Following isolation, RNA concentrations were normalized, and cDNA was synthesized using a high-capacity RNA to cDNA kit (ThermoFisher Scientific, 43874060), including a corresponding no RT condition for each synthesis reaction. RT-qPCR primers were designed using NCBI primer blast in combination with Ensembl and acquired from Thermo Fisher. Where possible, primers were designed to span exon-exon junctions. Primer GC contents ranged between 45-60% GC content and product sizes were between 70-179bp. Sequences for all primer pairs used are as follows: *Igf1:* fwd (5’ -> 3’): ACCTCAGACAGGCATTGTGG, rev: CGATAGGGACGGGGACTTCT*; Osm:* fwd: CAGCTGCAGAATCAGGCGAA, rev: GGTTTTGGAGGCGGATATAGGG*; Lifr:* fwd: CGTAGAAGAACTGGCTCCC, rev: CCTCGTCTTGGGCGTATCTC*; Osmr:* fwd: GGTCCTTCATCCAGCCTTCC, rev: GCTCCTCCAAGACTTCGCTT*; Il6st:* fwd: CGGCTCATATGGAAGGCACT, rev: CCCACCTTGTTTCTTGCTGC; *Tbp*: fwd: CTGGCGGTTTGGCTAGGTTT, rev: ACCATGAAATAGTGATGCTGGG. All primers were validated for product size and specificity via gel electrophoresis and visualization using cDNA from various brain region tissues including cortex, V-SVZ and cerebellum. All primers were within 90-110% efficiency, as confirmed by RT-qPCR derived efficiency curves calculated from serially diluted cDNA samples. Primers were diluted to 10 μM prior to use, with a final reaction concentration of 0.2 µM. RT-qPCR was performed using Fast SYBR™ Green Master Mix (Applied Biosystems, 4309155) in 20 μl reactions using an Applied Biosystems 7500 machine. Cycle times were: step 1: 50°C, 2 min (1 cycle), step 2: 95°C, 2 min (1 cycle) and, step 3: 95°C, 10 sec, 60°C, 30 sec (40 cycles).

Relative expression of each target gene was calculated via the ΔΔCt method using *Tbp* as a housekeeping gene which was tested against a range of housekeepers and selected based on based expression stability across experimental conditions. All RT-qPCR runs included blank and no RT controls.

### 10X Genomics Xenium tissue preparation and data processing

10X Xenium *in situ* based single cell spatial transcriptomics was performed on adult CD1 mouse brain tissue from vehicle-control or cuprizone-rapamycin demyelination with a three-week recovery period. Following RNAse-free removal of brains, fresh tissue was flash frozen in OCT embedding matrix and stored at -70°C. Subsequently, 10 µm cryosections were mounted onto Xenium slides (chemistry v1) following 10X Genomics guidelines. All tissues were equilibrated to -23°C in a cryostat prior to sectioning. Each section contained lateral ventricle and corpus callosum regions which were rostral of the hippocampus. One section was collected per animal. These slides were stored at -70°C prior to subsequent preparation steps.

Sectioned tissue was processed according to the Xenium workflow for fresh frozen tissue. To fix and permeabilize the tissue, sections were incubated for 1 minute at 37°C, then fixed in 4% PFA (Electron Microscopy Sciences) for 30 minutes at room temperature. Sections were washed in RNAse-free 1X PBS for one minute (Thermo Fisher), and subjected to a series of permeabilization steps and PBS washes. Permeabilization steps included 1% SDS incubation for 2 minutes and a chilled 70% methanol incubation for 1 hour. Following permeabilization, slides were washed twice with PBS and then placed in PBS/0.05% Tween-20 (PBS-T, Thermo Fisher). Sections were hybridized with probe solution, containing a 347 gene probe panel (see Suppl. Table 2) in TE-buffer, for 22 hours at 50°C. Afterwards, sections underwent several PBS washes and were incubated at 37°C in post-hybridization wash buffer. Following a series of PBS-T washes, sections were incubated in Xenium ligation enzymes for 2 hours at 37°C. Following multiple rounds of PBS-T washes, probe amplification was achieved by incubating sections for 2 hours at 30°C in Xenium amplification enzyme solution. Sections were washed twice in TE buffer, and stored overnight at 4°C.

Autofluorescence quenching and nuclei staining were next performed. Sections were washed in PBS, incubated in reducing agent for 10 minutes, washed in 70% and 100% ethanol, and incubated in Xenium autofluorescence solution for 10 minutes. Sections were then washed three times in 100% ethanol, dried at 37°C for 5 minutes and rehydrated via a series of PBS and PBS-T washes. Nuclear staining buffer was added to sections for 1 minute and sections washed 4 times in PBS-T. Samples were loaded into the Xenium analyzer instrument (software v1.4.0.6) and subjected to several cycles of reagent application, probe hybridization, imaging, and probe removal. Pre-processing of captured Z-stack images was performed using the Xenium on-board analysis pipeline^31^. In brief, a custom Xenium codebook (described below) was used to decode imaged fluorescent puncta into transcripts, with each decoded fluorescence signature assigned a codeword which was associated to a target gene. Quality scores (Q-Scores) were assigned to transcripts based on maximum likelihood codewords compared to the likelihood of other sub-optimal codewords, to provide the confidence in each decoded transcripts assigned identity. Only transcripts with a Q-score ≥ 20 were included in downstream analysis. Negative control codewords (codewords that do not correspond to any probe) and negative control probes (probes included in the panel which do not match any biological sequence) were utilized to assess decoding accuracy and assay specificity, respectively. On-instrument cell segmentation was performed based on 3D DAPI morphology. Nuclei identification and cell boundaries were subsequently flattened into a 2D mask, cell IDs allocated to each identified cell, and transcripts assigned to cell IDs based on their X-Y co-ordinates. The Xenium on-board analysis default boundary for expansion (15 µm) was set for all sections.

### 10X Xenium data analysis

Standardized Xenium output files were exported for downstream analyses. Data were visualized in Xenium Explorer (version 1.3.0), with regions of interest (ROIs) defined using the freehand select tool and cell IDs exported. Transcript count, feature, coordinate and cell ID data were imported to R and Seurat objects created (v5.0.1) via the LoadXenium function. Subsets were then produced from object data using the previously exported cell IDs to include only cells contained within our ROI, and these regions were concatenated where multiple ROIs were specified on one section. All downstream Xenium analysis was conducted in R package Seurat. Initial quality checks assessed cells based on the number of detected genes and total transcript counts per cell. Low quality cells with greater than +/-2.5 standard deviations from the mean in either of these parameters were excluded, with an average exclusion of approximately 4% of cells per dataset.

Filtered data were normalized using SCTransform^85^, and PCA based on highly variable genes performed to reduce dimensionality and capture the main sources of variation in the data. A shared nearest neighbor (SNN) graph was constructed using Seurat FindNeighbors based on the principal components identified through PCA. A Louvain clustering algorithm implemented in Seurat (FindClusters) was applied to partition cells into distinct clusters based on their transcriptomic profiles. The most conservative, yet biologically meaningful, clustering resolution parameter was chosen.

UMAPs and image dimensional reduction plots visualizing cell co-ordinates (ImageDimPlot) were employed to visualize the data in two-dimensional space. Seurat FeaturePlots, ImageFeaturePlots and heatmaps were used to investigate gene expression, annotate and interpret cell clustering patterns. We used the following well-defined markers genes to identify and annotate cell types and states: For NSCs (both activated and dormant), *Nes, Veph1,Vnn1, Notum, Tspan18*, *Egfr, Ascl1*; for TAPs, *Mki67, Rrm2, Hells, Gsx2*; for astrocytes, *Aqp4, Agt, Hsd11b1, Lcat*; for all oligolineage cells: *Olig1, Sox10*; for immature oligodendrocytes, *Enpp4*; for OPCs, *Pdgfra, Cspg4*; for mature oligodendrocytes, *Mog, Mag, Opalin;* for all neuroblasts, *Dlx1, Dlx2, Sp8, Sp9, Gad1, Gad2*; for all microglia, *Aif1, Trem2, Cx3cr1, Tmem119*; for other immune cells including T cells and neutrophils, *Cd52*, *Cd69*; for striatal neurons, *Calb1, Gad1, Gad2, Bcl11b*; for endothelial cells, *Pecam1, Esam, Plvap*; for pericytes, *Carmn, Cspg4, Ano1*; for VAMCs, *Pdgfrb, Myh11, and Mylk*; and for ependymal cells, *Foxj1, Pvalb*. Annotation of single cell spatial transcriptomics data was also informed by prior knowledge of expected cell location.

Each Xenium field of view (FOV) was processed and annotated individually and Seurat SelectIntegrationFeatures, FindIntegrationAnchors and IntegrateData functions used to merge FOVs. Where relevant for improved annotation, further subsets of data were created using Seurat subset function. Normalization, PCA, SNN analysis and clustering were performed on merged datasets and/or subsets as described above.

### MERSCOPE multiplexed error-robust fluorescent *in situ* hybridization (MERFISH) tissue preparation and data processing

MERFISH (as described by ^34–36^) single cell spatial transcriptomics was performed using tissue from CD1 adult mice which had undergone either control-vehicle or cuprizone-rapamycin demyelination with a three-week recovery period. Under RNase-free conditions, brains were removed and flash frozen in OCT embedding matrix. Tissues were stored at -70°C until further processing. Tissues were equilibrated in a -23°C cryostat for at least one hour prior to sectioning and one 10 µm cryosection from each brain was collected on a MERSCOPE slide (20400001). Each slide was stored in an individual 60 mm petri dish throughout the tissue preparation process.

Sections were processed in accordance with MERSCOPE guidelines for fresh frozen tissue (protocol no. 91600002), beginning with fixation and permeabilization. All incubations were performed stationary at room temperature in 5 mL of solution, unless otherwise stated. In brief, sections were washed 3 times with 1X PBS and incubated in 70% ethanol overnight at 4°C. Following a PBS wash, a small volume of blocking solution (cell boundary blocking buffer premix (MERSCOPE, 20300100) containing 5% RNase inhibitor (NEB, M0314L)) was added directly to the sections. Sections were covered with a piece of parafilm and incubated for an hour. Following parafilm removal and blocking buffer aspiration, MERSCOPE primary staining solution (5% RNase inhibitor and 1% cell boundary primary stain mix (MERSCOPE, 20300010) in cell boundary blocking buffer premix) was added in a small volume directly on top of the sections. Sections were covered again and allowed to incubate for 1 hour. Sections were washed with PBS 3 times with agitation, and secondary staining solution (5%RNase inhibitor, 3% cell boundary secondary stain mix (MERSCOPE, 20300011) in cell boundary blocking buffer premix) was added. Sections were covered with parafilm and incubated for 1 hour. Following a further 3 PBS washes with agitation, sections were incubated in fixation buffer (4% paraformaldehyde in 1X PBS) for 15 minutes. Tissues were washed two more times with PBS and once with sample prep wash buffer (MERSCOPE, 20300001) before proceeding to encoding probe hybridization.

Sections were incubated in formamide wash buffer (MERSCOPE, 20300002) for 30 minutes at 37°C. After careful aspiration, the custom MERSCOPE gene probe panel (see Suppl. Table 2) was added directly on top of the tissue. Parafilm was added to cover sections and petri dishes were sterilized, sealed and placed in a humidified incubator (37°C) for 36-48 hours. Subsequent to encoding probe hybridization, sections were washed twice with formamide wash buffer (47°C, 30 minutes per wash) and once with sample prep wash buffer for 2 minutes. Sections were gel embedded using the MERSCOPE gel embedding solution (20% gel embedding premix (MERSCOPE, 20300004), 10% N,N,N’,N’-tetramethylethylenediamine in 10% w/v ammonium persulfate solution). Solution was added directly to tissue, covered with a gel slick (VWR, 12001-812) coated glass coverslip and allowed to set for a minimum of 1.5 hours. Once completely set, coverslips were carefully removed and pre-warmed (30°C) clearing solution (MERSCOPE, 20300003) supplemented with 1% proteinase K (NEB, P8107S) was added. Tissues were incubated in clearing solution for a minimum of 24 hours and a maximum of 4 days prior to DAPI and PolyT counterstaining with the MERSCOPE DAPI and PolyT Staining Reagent kit. Where required, the clearing buffer was replenished after 3 days. Activated imaging cartridges and samples were loaded into the MERSCOPE instrument in accordance with the Instrument User Guide (protocol no. 91600001). We used a custom gene panel codebook, containing the unique, binary barcodes for each targeted gene. This panel codebook contained 15 ‘blank’ barcodes which did not match any target gene to be used in instances where barcode matches or nearest correct barcode assignments could not be made with confidence. To generate binary barcodes for each detected transcript, samples underwent multiple, sequential rounds of fluorescent readout probe hybridization, field of view imaging and fluorophore extinguishing. After imaging, data were decoded to produce a binary barcode based on the detection (or absence) of readout probes in each imaging round. Generated barcodes were assigned to appropriate target genes based on barcode matches or, in the case of readout errors, their nearest correct barcode or a blank barcode. Cell segmentation was also performed on the instrument using the Cellpose cell segmentation algorithm^86^ which accounted for DAPI, PolyT and cell boundary labelling images obtained by the instrument.

### MERFISH data analysis

Standardized MERSCOPE output files were exported including gene expression, cell metadata, co-ordinate and cell boundary information. Data for each section were first visualized using MERSCOPE Vizualizer software. V-SVZ and adjacent corpus callosum ROIs were outlined using the polygonal selection tool and cell ID-containing HDF5 files were exported for downstream analysis. All data were processed using python (v3.8.0). Previously exported HDF5 files were used to extract and process only our specified ROI. All transcripts assigned to blank barcodes were removed. For each section, median cell volumes were calculated and cells with volumes 3 times higher than the median were removed. Scrublet ^87^ was then used to predict and remove potential doublets based on the transcript count matrix. For data with a binomial distribution, doublet score thresholds were set to the mid-point between distribution peaks. If data did not follow a binomial distribution, thresholds were set to the distribution curve plateau. This initial filtering excluded an average of around 9% of cells per dataset.

Data were then loaded into Seurat in R using the ReadVizgen function. Additional inputs were loaded, including cell centroid and segmentation information, using the CreateCentroids and CreateFOV functions in Seurat. Seurat objects containing transcript, cell ID, co-ordinate and segmentation data were via CreateSeuratObject. In a similar manner as performed for Xenium data (see *10X Xenium data analysis*), data were normalised using SCTransform, PCA was performed using highly variable genes and an appropriate number of principle components selected using the Elbow method. SNN graphs were constructed using Seurat FindNeighbours, and FindClusters was used to cluster cells based on their gene expression. As with Xenium data, UMAPs were created using Seurat RunUMAP to reduce dimensionality and visualize data. Seurat ImageDimPlot was used to visualise the co-ordinates and cluster information for each cell.

MERFISH data required a further round of filtering. Cells greater than +/-2.5 standard deviations from mean transcript counts or mean genes expressed per cell were excluded. Cells were also excluded where cell-type annotation for a particular cluster was not possible, even at high clustering resolutions, due to ambiguous marker gene expression. Across all MERFISH datasets, this additional filtering of low quality and ambiguous cells excluded approximately 33% of cells. The higher rate of exclusion likely reflects cell segmentation inaccuracies in which more doublets, cellular fragments and non-cells were identified by MERFISH, as compared to Xenium datasets.

As previously described, FeaturePlots, ImageFeaturePlots and heatmaps were employed to visualize gene expression, annotate and interpret cell clustering. We used the following well-defined markers genes to identify and annotate cell types and states: For NSCs (activated and dormant), *Nes, Veph1,Vnn1, Notum, Tspan18*, *Egfr, Ascl1*; for TAPs, *Mki67, Rrm2, Hells, Gsx2*; for astrocytes, *Aqp4, Agt, Hsd11b1, Lcat*; for all oligolineage cells: *Olig1, Sox10*; for immature oligodendrocytes, *Enpp4*; for OPCs, *Pdgfra, Cspg4*; for mature oligodendrocytes, *Mog, Mag, Opalin;* for all neuroblasts, *Dlx1, Dlx2, Sp8, Sp9, Gad1, Gad2*; for all microglia, *Aif1, Trem2, Cx3cr1, Tmem119*; for other immune cells including T cells and neutophils, *Cd52*, *Cd69*; for striatal neurons, *Calb1, Gad1, Gad2, Bcl11b*; for endothelial cells, *Pecam1, Esam, Plvap*; for pericytes, *Carmn, Cspg4, Ano1*; for VAMCs, *Pdgfrb, Myh11, and Mylk*; and for ependymal cells, *Foxj1, Pvalb*. We also informed our annotation of spatial transcriptomics data with the spatial location of cells.

Resultant individual Seurat objects for each MERFISH section were merged as previously described for Xenium data, using Seurat SelectIntegrationFeatures, FindIntegrationAnchors and IntegrateData functions. Further subsets of data were created where required for improved annotation and analysis using Seurat subset. Normalization, PCA, SNN analysis and clustering were re-run on merged datasets and/or subsets as described in previous sections.

### Probe panel design for *in situ*-based single cell spatial transcriptomics

Our Xenium experiments used a 347-probe panel comprised of 247 probes from the 10X Genomics pre-designed mouse brain panel and 100 custom add-on probes which were designed to include key marker genes, ligands and receptors. The number of probes included for each target gene was optimized by Xenium’s probe panel designer to reduce the likelihood of optical crowding based on relevant adult CD1 control and cuprizone-rapamycin datasets. MERFISH experiments used a 300-probe fully custom panel, consisting of marker genes, ligands and receptors. This panel was designed using the MERSCOPE online panel designer and probe selections were informed by prior assessment of gene specificity and expression level in our relevant scRNA-seq datasets. Further information, including all custom genes included in each panel, can be found in Suppl. Table 2.

### Cortical precursor cell cultures

Cortical precursor cultures were prepared as previously described^53,88^. Briefly, for each culture, E15-16 CD1 embryos of both sexes were collected from the same mother. Meninges were removed and cortices exposed and placed in ice cold Hank’s buffered salt solution (HBSS). Cortical tissue directly above the ventricle was collected and mechanically triturated to dissociate cells. Dissociated cortical precursor cells were plated at a density of 150,000 cells/well on 2% laminin (Corning)/1% poly-D-lysine (Sigma) glass coverslips precoated 24 hours prior. Cultures were maintained at 37°C in the Neurobasal medium (GIBCO) supplemented with 2% B27 (Life Technologies), 0.5 mM L-glutamine (Lonza), 15 ng/mL FGF2 (Corning), and 1% penicillin/streptomycin (Lonza), except for those included in the ‘high’ FGF concentration group (Figure 6A). This latter group were instead maintained in Neurobasal medium (GIBCO) supplemented with 2% B27 (Life Technologies), 0.5 mM L-glutamine (Lonza), 50 ng/mL FGF2 (Corning), and 1% penicillin/streptomycin (Lonza).

### EdU treatment of cortical precursor cell cultures

EdU (5-ethynyl-2’-deoxyuridine) incorporation was used to measure proliferation of cultured cortical precursor cells. EdU assays were performed using the Click-iT™ EdU Cell Proliferation Kit (Thermo Fisher, C10337). In brief, cortical precursor cultures were prepared as described above, in media supplemented with either 15 ng/mL or 50 ng/mL FGF2 (see *Cortical precursor cell cultures*) and maintained for 3 days in vitro (DIV). At 3 DIV, 1µM of EdU was added and cells were incubated for 24 hours. Cells were fixed with 4% PFA for 10 minutes, washed 3 times with 1X PBS and permeabilized with 0.3%Triton-X100 in PBS for 30 minutes at room temperature. EdU was detected and labelled with AlexaFluor-488 dye by incubating cells in the Click-iT reaction buffer (prepared according the manufacturer protocol) for 30 minutes at room temperature in the dark. Following 3 further PBS washes, cells were incubated in DAPI (1:1000) for 5 minutes to label nuclei. Cells were washed, coverslips mounted with Vectashield (CedarLane, VECTH1000) and sealed prior to imaging.

### Ligand treatment of cortical precursor cell cultures

Cortical precursor cell cultures were treated with either vehicle (0.1% BSA in PBS), IGF-1 (100 ng/ml) or OSM (100 ng/ml) to assay their effects on precursor proliferation and oligodendrogenesis. Cultures were treated with either ligand or vehicle at 4 hours post-plating and at 3 DIV. At 4 DIV, cells were prepared for subsequent immunostaining.

### Antibodies

We used the following antibodies for our immunostaining studies: Goat anti-PDGFRα (1:250, R&D Systems), rabbit anti-OLIG2 (Abcam), mouse anti-CC1 (1:100, Calbiochem), Rabbit anti-Ki67 (1:200 immunohistochemistry, 1:300 immunocytochemistry, Abcam), Rat anti-SOX2 (1:300, eBioscience), anti-goat Alexa555 (1:1000, Invitrogen), anti-rat Alexa488 (1:1000, Invitrogen), anti-rabbit Alexa647 (1:1000, Invitrogen), anti-mouse Alexa647 (1:1000, Jackson ImmunoResearch Laboratories).

### Immunocytochemistry on cortical precursor cultures

To assay proliferation and oligodendrogenesis, 4 DIV cortical precursor cells were immunostained for Ki67 (1:200) and PDGFRα (1:250), respectively. Culture media was removed and cells fixed in 4% PFA at room temperature for 10 minutes. Cells were then washed 3 times in 1X PBS, and blocked and permeabilized with 0.3% Triton-X100 and 5% BSA in PBS for 1 hour at room temperature. Primary antibodies, PDGFRα (1:250) and Ki67 (1:300), were diluted in the aforementioned blocking/permeabilization buffer and added directly to coverslips in a humidified chamber. Cells were incubated in primary antibodies overnight at 4°C, washed 3 times with PBS and secondary antibodies added (for Ki67: anti-rabbit Alexa647, for PDGFRα: anti-goat Alexa555, also see *Antibodies* section). Secondary antibodies were diluted in 5% BSA-PBS and incubated with cells for 1 hour at room temperature in the dark. Secondary antibodies were washed from cells (3 times with PBS). To counterstain nuclei, DAPI (1:1000, diluted in PBS) was incubated with cells for 5 minutes in the dark at room temperature. A further 3 PBS washes were performed before coverslips were mounted to slides with Vectashield (Vectorlabs) mounting medium and sealed prior to imaging.

### Immunohistochemistry on adult tissue sections

For *NestinCreErt2-Eyfp* lineage tracing and *in vivo* ligand infusion experiments, mice were perfused with ice-cold PBS and 4% PFA and brains collected. Brains were immersion fixed in 4% PFA overnight and cryopreserved via a sucrose gradient (15% sucrose, then 30% sucrose in PBS). Following immersion in 30% sucrose for 24 hours, brains were flash frozen in OCT embedding matrix and stored at -70°C prior to sectioning. Brains were equilibrated to -23°C in a cryostat and 14 µm coronal sections were collected across the rostral-caudal axis of the lateral ventricles. Sections were stored at -70°C prior to further processing. Immunohistochemistry was performed on sections, with slides from each condition processed at the same time to reduce batch variability. Sections were dried for 30 minutes at 37°C, briefly rehydrated in 1X PBS, and blocked and permeabilized for 1 hour at room temperature in 0.3% Triton-X100 and 5% BSA PBS. For primary antibodies raised in mouse, a MOM (mouse-on-mouse) kit was used according to the manufacturer’s protocol (Vector Laboratories). Sections were incubated with relevant primary antibodies at their optimized concentrations (see *Antibodies* section) overnight in a humidified environment at 4°C. Sections were washed 3 times with PBS prior to application of appropriate secondary antibodies at optimized concentrations (see *Antibodies* section). Following a 1-hour room temperature incubation in the dark, sections were washed 3 times with PBS, counterstained with DAPI (1:1000) for 5 minutes at room temperature, and washed a further 3 times with PBS. Coverglasses were applied to tissue with Vectashield and sealed.

### Imaging of immunofluorescent staining

For cortical precursor culture immunocytochemistry, a Zeiss Axio Imager M2 system with an X-Cite 120 LED light source and C11440 Hamamatsu camera was used to obtain stitched images. Cells were imaged with a 20X objective. Four quadrants of equal size were imaged for each sample (e.g. four scans per coverslip). Quadrants were selected to avoid coverslip edges, but were otherwise randomly chosen. V-SVZ images from immunostained brain tissue sections were collected using a Zeiss Axio Imager M2 system with an X-Cite 120 LED light source, Apotome 3 and and C11440 Hamamatsu camera (for image analysis), or a Zeiss LSM900 with an Airyscan 2 detector (for representative images). For *NestinCreErt2-Eyfp* mice, 3 anatomically matched sections from each rostral and caudal aspect were imaged per sample. For ligand infusion experiments, two stitched images containing the V-SVZ dorsal and lateral walls, adjacent corpus callosum, and nearby striatum were collected for each sample.

## QUANTIFICATION AND STATISTICAL ANALYSIS

### Quantification of immunofluorescence data

For lineage tracing morphological analysis, marker-positive cells were counted from imaged rostral and caudal corpus callosum regions, including double- and triple-positive cells. Results are presented as either the quantification of one EYFP-positive cell type (e.g. OLIG2+, PDGFRa+, EYFP+ OPCs) as a percentage of the EYFP-negative cells for the same cell type (e.g. OLIG2+, PDGFRa+, EYFP-OPCs), or expressed as density relative to the quantified corpus callosum area.

For cortical precursor experiments assessing the effects of FGF2, IGF-1 and OSM, all normal nuclei, PDGFRα-positive, Ki67-positive and PDGFRα, Ki67-double positive cells were counted. Results are presented quantifying total cell number, PDFGFRα-positive cells as a percentage of all cells, Ki67-positive cells as a percentage of all cells, Ki67, PDGFRα-double positive cells as a percentage of all PDGFRα-negative cells for each ligand and its respective control. A minimum of 200 total cells were quantified per image. For experiments quantifying EdU-incorporation and cell death, apoptotic cells were defined as having a small, compacted, relatively circular morphology with intense labelling comparatively to other DAPI stained nuclei. These were not included in the quantification of EdU-positive nuclei. An average of 1490 total nuclei were quantified for each independent sample.

For ligand infusion experiments, marker positive and double-positive cells were counted for cells at the V-SVZ lateral wall, dorsal wall and corpus callosum. For quantification, cells included in V-SVZ dorsal wall and corpus callosum were pooled. Dorsal wall/corpus callosum and lateral wall quantification results are reported for both ligands and their respective controls as total SOX2-positive PDGFRα-negative cells, total PDGFRα-positive cells, total Ki67-positive cells, total Ki67-positive SOX2-positive cells, and total Ki67-positive PDGFRα-positive cells.

### Quantification of cell proportions in scRNA-seq datasets

The proportion of microglia reported in Figure 1H were quantified by extracting the number of microglia (as annotated by well-defined cell type markers) in CD1 and *NestinCreErt2-Eyfp* control-vehicle and cuprizone-rapamycin datasets. The number of microglia were divided by the total number of cells for each respective dataset and expressed as a percentage. The proportions of each neural cell type as a percentage of total neural cells (reported in Figure 1K) were quantified by extracting the number of cells of each cell type (as annotated by defined cell-type markers) from CD1 and *NestinCreErt2-Eyfp* vehicle-control and cuprizone-rapamycin neural subsets (Figure 1I). The number of cells of each neural cell type was divided by the total number of neural cells for each respective dataset and expressed as a percentage. For quantification of *Eyfp* expression reported in Figure 2E, the percentage of *Eyfp*-positive and negative cells were computed for each cell type in vehicle-control and cuprizone-rapamycin *NestinCreErt2-Eyfp* datasets. The percentage of *Eyfp*-positive cells for each cell type was reported for each dataset. Neural precursor cell proportions were quantified using the neural precursor subset shown in Figure 2F. The number of dNSCs, aNSCs and TAPs was quantified from each dataset of origin, then divided by the total number of neural precursors in their respective dataset and expressed as a percentage. To quantify the proportion of proliferating TAPs, the number of TAPs in G2, M, and S phases (as determined by the Cyclone method) were counted and divided by the total number of TAPs in each dataset.

### Pearson correlation analysis

Comparisons of gene expression between dNSCs, aNSCs and TAPs from CD1 control and cuprizone rapamycin datasets (Figure 2I and Suppl. Figure 2I) were performed by averaging the expression of each gene across all cells. The Pearson correlation coefficient was then determined by using the cor.test function in R. Pearson correlation coefficients (*r*) values are shown in the corresponding figure and figure legend for each comparison.

### Quantification of Xenium data

The proportions of cells of the oligodendrogenic lineage, including OPCs, immature and mature oligodendrocytes in the dorsal V-SVZ and corpus callosum were quantified for each individual Xenium dataset (two animals each for vehicle and cuprizone-rapamycin). To this end, ROIs containing only the dorsal wall and adjacent corpus callosum were extracted and counts of OPCs, immature and mature oligodendrocytes, total oligodendrocyte lineage cells and total cells were quantified from spatial plots using Adobe Illustrator software. Data for each cell type were expressed as percentages of total oligolineage cells and total cells for each Xenium dataset.

### Statistical analysis

Sample sizes (N) for each experiment are noted in associated figure legends. For scRNA-seq data, N refers to the number of independent scRNA-seq preparations and runs for each condition. For cortical precursor cultures, N refers to the number of independent cultures from embryos derived from different litters for each condition. For Xenium and MERFISH, N refers to the number of independent sections from different animals for each condition. For immunhistochemical analysis, N refers to the number of brains analyzed for each condition. Details of statistical analysis relevant to each experiment are contained within associated legends. For two group quantification comparisons (e.g. vehicle control vs. cuprizone rapamycin, control vs. IGF-1, control vs. OSM), normality assumptions were confirmed and independent two-tailed Student’s t-tests performed. These analyses were performed using GraphPadPrism8 software, and results considered statistically significant if p<0.05. In all figures, asterisks denote statistical significance where *p < 0.05, **p < 0.01 and ***p<0.001.

## SUPPLEMENTAL FIGURE LEGENDS

**Supplemental Figure 1. L*i*neage *tracing and scRNA-seq to characterize oligodendrogenesis in the control and remyelinating V-SVZ and corpus callosum.*** Related to Figure 1. (A) Low magnification image of the top of the dorsal V-SVZ and corpus callosum in a coronal forebrain section from an adult *NestinCreErt2-Eyfp* mouse treated with tamoxifen for five consecutive days and analyzed immediately for EYFP (green) and counterstained with DAPI (white) to show nuclei. The boxed region is shown at higher magnification as an inset. Note that cells in the V-SVZ and not corpus callosum were labelled. Scale bars = 100 µm and 50 µm (for insets). **(B)** Representative confocal images of the dorsal V-SVZ and corpus callosum of a control adult *NestinCreErt2-Eyfp* mouse eight weeks post-tamoxifen, analyzed for EYFP and immunostained for OLIG2 (red) and PDGFRα (blue). The boxed region is shown at higher magnification in the bottom panels. The arrow denotes a triple-labeled NPC-derived OPC and the arrowhead an EYFP-positive, OLIG2-positive, PDGFRα-negative NPC-derived oligodendrocyte. Scale bar = 100 um. **(C-H)** Six-week-old CD1 mice were treated with cuprizone-rapamycin or vehicle for six weeks, and dorsal and lateral V-SVZ tissue including the corpus callosum was collected for scRNA-seq. (**C, D, F, G**) t-SNE cluster map visualizations of all V-SVZ transcriptomes in these datasets, annotated for cell types. (C and D) are preparations of control tissue collected either immediately after (0 week) or at 3 weeks post-treatment, while (F and G) are preparations of cuprizone-rapamycin tissue isolated 3 weeks post-treatment. Transcriptionally-distinct clusters are colour-coded, and the plot is annotated for cell types within these clusters, as determined by analysis of well-characterized marker genes. dNSC = dormant NSCs, aNSC = activated NSCs, TAPs = transit amplifying precursors, OLGs = oligodendrocytes. VAM = vasculature-associated mesenchymal cells. The control 0 weeks recovery dataset (C) contained 4448 cell transcriptomes and was analyzed at resolution 0.8 with 22 identified clusters. The control with 3 weeks recovery dataset (D) contained 4127 cell transcriptomes and was analyzed at resolution 1.2 with 21 identified clusters. The cuprizone-rapamycin datasets (F, G) contained 3675 and 2573 cell transcriptomes, respectively. The first cuprizone-rapamycin dataset (F) was analyzed at resolution 1.6 with 24 identified clusters, and the second (G) was analyzed at resolution 1.6 with 23 identified clusters. **(E, H)** t-SNE visualizations of the datasets in (D) and (G), respectively, overlaid for expression of marker genes that are specific to the different cell types in these datasets. Expression levels are coded as per the adjacent keys. **(I-K)** Six-week-old *NestinCreErt2-Eyfp* mice were injected with tamoxifen at 5 weeks, then treated with vehicle for six weeks, and three weeks post-treatment the dorsal and lateral V-SVZ tissue including the corpus callosum was collected and scRNA-seq was performed. (I and J) show t-SNE cluster map visualizations of all V-SVZ transcriptomes in these datasets annotated for cell types and (K) shows the dataset in (J) overlaid for expression of marker genes specific to different cell types. Expression levels are coded as per the adjacent keys. The first *NestinCreErt2-Eyfp* control dataset (I) contained 5508 cell transcriptomes and was analyzed at resolution 0.4 with 15 identified clusters. The second *NestinCreErt2-Eyfp* control dataset (J) contained 4211 cell transcriptomes and was analyzed at resolution 1.2 with 21 identified clusters.

**Supplemental Figure 2. s*c*RNA*-seq analysis of V-SVZ and corpus callosum neural cells and NPCs*.** Related to Figures 1 and 2. (A-C) Six-week-old *NestinCreErt2-Eyfp* mice were injected with tamoxifen at 5 weeks, then treated with cuprizone-rapamycin for six weeks, and three weeks post-treatment the dorsal and lateral V-SVZ tissue including the corpus callosum was collected for scRNA-seq. (A and B) show t-SNE cluster visualizations of all V-SVZ transcriptomes in these datasets annotated for cell types and (C) shows the dataset in (B) overlaid for expression of marker genes specific to different cell types. Expression levels are coded as per the adjacent keys. NSCs = neural stem cells, TAPs = transit amplifying progenitors, OLGs = oligodendrocytes. VAM = vasculature-associated mesenchymal cells. The first *NestinCreErt2-Eyfp* cuprizone-rapamycin dataset (A) contained 4324 cell transcriptomes and was analyzed at resolution 0.4 with 15 identified clusters. The second *NestinCreErt2-Eyfp* cuprizone-rapamycin dataset (B) contained 11030 cell transcriptomes and was analyzed at resolution 0.8 with 26 identified clusters. **(D)** Neural cells were subsetted from the 8 individual control and cuprizone-rapamycin treated datasets in Suppl. Figures 1 and 2, merged and reanalyzed. Both top and bottom panels show the merged neural cell datasets color-coded for whether they were from the CD1 or *NestinCreErt2-Eyfp* datasets. The top panel shows the merged dataset before Harmony-mediated batch correction and the bottom after one round of batch correction. **(E, F)** t-SNE visualizations showing the merged, batch-corrected total neural cell dataset from (D). In (E) the cluster diagram is annotated for cell types and in (F) it is overlaid for expression of cell type-specific marker genes. Expression levels in (F) are coded as per the adjacent keys. **(G, H)** dNSCs, aNSCs and TAPs were subsetted from the total neural cell dataset in (E) and reanalyzed through the pipeline. (G) shows the t-SNE cluster visualization, annotated for cell types, and (H) shows expression overlays for marker genes that distinguish the different precursor types.

Expression levels are coded as per the adjacent keys. **(I)** Pearson correlation analysis of each detected gene in dNSCs from the dataset shown in (Fig. 2F), comparing averaged gene expression of control (y-axis) and cuprizone-rapamycin treated (x-axis) transcriptomes.

**Supplemental Figure 3. C*h*aracterization *of microglia in scRNA-seq datasets, and of all cells in the V-SVZ and corpus callosum in Xenium datasets*.** Related to Figures 3 and 4. (A, B) t-SNE visualizations of the merged total cell CD1 V-SVZ datasets (two control and two cuprizone-rapamycin treated) showing (A) the cluster diagram annotated for cell types, and (B) color-coded for their treatment condition. The merged CD1 control and cuprizone rapamycin dataset contained 14832 cell transcriptomes and was analyzed at resolution resolution 1.2 with 31 identified clusters. **(C, D)** t-SNE visualizations of microglia from the dataset shown in (A, B), subsetted and reanalyzed. (C) shows the cluster diagram with the two groups of transcriptionally-distinct microglia outlined, while (D) shows expression overlays for pan-microglial genes (*Aif1, Cx3cr1, Trem2*) and genes more highly expressed in the remyelination-enriched microglia (*Apoc4, Cybb, Lgals3*) or in homeostatic microglia (*Bank1, Crybb1, Il7r, Ecscr*). Expression levels are coded as per the adjacent keys. **(E-G)** Coronal sections from vehicle or cuprizone-rapamycin treated mice were obtained 3 weeks post-treatment and analyzed by Xenium-based single cell multiplexed *in situ* hybridization with a probeset targeting 347 genes (in Suppl. Table). The ROI was defined as the region around the V-SVZ plus the corpus callosum. Cellular transcriptomes within the ROI of an individual section were analyzed after removal of poor-quality transcriptomes. Transcriptomes from two control sections from two separate animals (E) or two remyelinating sections from two separate animals (F) were merged. The datasets in (E and F) were then merged together (G; also in Fig. 3F, G). Shown are the UMAP cluster visualizations, annotated for cell types. Each dot represents a single transcriptome. VAMC – vasculature-associated mesenchymal cell, OLGs = oligodendrocytes. The merged control Xenium dataset contained 14490 cells and was analyzed at resolution 2.4 with 31 identified clusters. The merged cuprizone-rapamycin Xenium dataset contained 16309 cells and was analyzed at resolution 2.8 with 36 identified clusters. (H) UMAP visualization of the merged dataset in (G) overlaid for expression of cell type-specific marker genes. Expression levels are coded as per the adjacent keys.

**Supplemental Figure 4. X*e*nium *and Merscope-based single cell transcriptomics to analyze the control and remyelinating V-SVZ and corpus callosum*.** Related to Figures 3 and 4. (A, B) UMAP cluster visualizations of astrocytes, ependymal cells, NSCs and TAPs that were subsetted from the Xenium merged dataset shown in Fig. 3F and reanalyzed. In (A) the UMAP is annotated for cell types, while (B) shows expression overlays for genes characteristic of each of the cell types. Expression levels are coded as per the adjacent keys. This subset contained 8520 cells and was analyzed at resolution 1.2 with 19 identified clusters. **(C)** Spatial plot of the ROI in one representative section each of the control and remyelinating V-SVZ and corpus callosum showing ependymal cells (grey), endothelial cells (orange) and pericytes (red), as analyzed by Xenium. **(D)** The top panels show gene expression overlays for mRNAs that are differentially expressed in homeostatic versus remyelination-enriched microglia, shown on the enlarged microglial clusters (1 and 20) from the annotated UMAP in Figure 3F. The bottom panel shows a spatial plot for *H2-Eb1* mRNA expression in the ROI of representative control and remyelinating sections, as determined using Xenium. Relative expression levels are coded as per the adjacent keys. **(E-G)** UMAP cluster visualizations of merged control and cuprizone-rapamycin treated transcriptomes, as determined using MERFISH and shown in Fig. 4J. (E) shows the UMAP with the transcriptomes color-coded for their condition of origin. The merged MERFISH dataset contained 19531 cells and was analyzed at resolution 2.0 with 22 identified clusters. (F and G) show the UMAP (F) annotated for cell types and (G) overlaid for the expression of cell type-specific marker genes. Expression levels are coded as per the adjacent keys. **(H)** Gene expression overlays for mRNAs that are differentially expressed in homeostatic versus remyelination-enriched microglia, shown on the enlarged microglial clusters (18 and 14) from the annotated MERFISH UMAP in panel (F).

**Supplemental Figure 5. *Analysis of the lateral V-SVZ after infusion of IGF1 or OSM.* Related to Figures 6 and 7. (A)** t-SNE visualizations of the neural cells in a previously-published scRNA-seq dataset (Borrett et al., 2020; GEO: GSE152281) from the postnatal day 6/7 (P6-P7) V-SVZ. The left panel shows the cluster visualization, annotated for cell types, and the right panels show expression overlays for mRNAs encoding IGF1 and OSM receptors. Expression levels are coded as per the adjacent keys. **(B)** Schematic of the growth factor infusion protocol. **(C)** Representative high magnification confocal images of the lateral V-SVZ of mice infused with OSM or 0.1%BSA/PBS (Control) and immunostained for Sox2 (yellow), PDGFRα (blue), and Ki67 (magenta). Also shown is DAPI counterstaining to highlight nuclei (white). The boxed regions are shown at higher magnification in the bottom panels. Arrows, filled arrowheads and empty arrowheads denote cells positive for only PDGFRα, PDGFRα and Ki67, and Sox2 and Ki67, respectively. Scale bars = 100 µm (tiled image) and 50 µm (insets).

