## Supplementary figures and images for "Single cell approaches define forebrain neural stem cell niches and identify microglial ligands that enhance precursor-mediated remyelination"

### Supplemental Figure 1

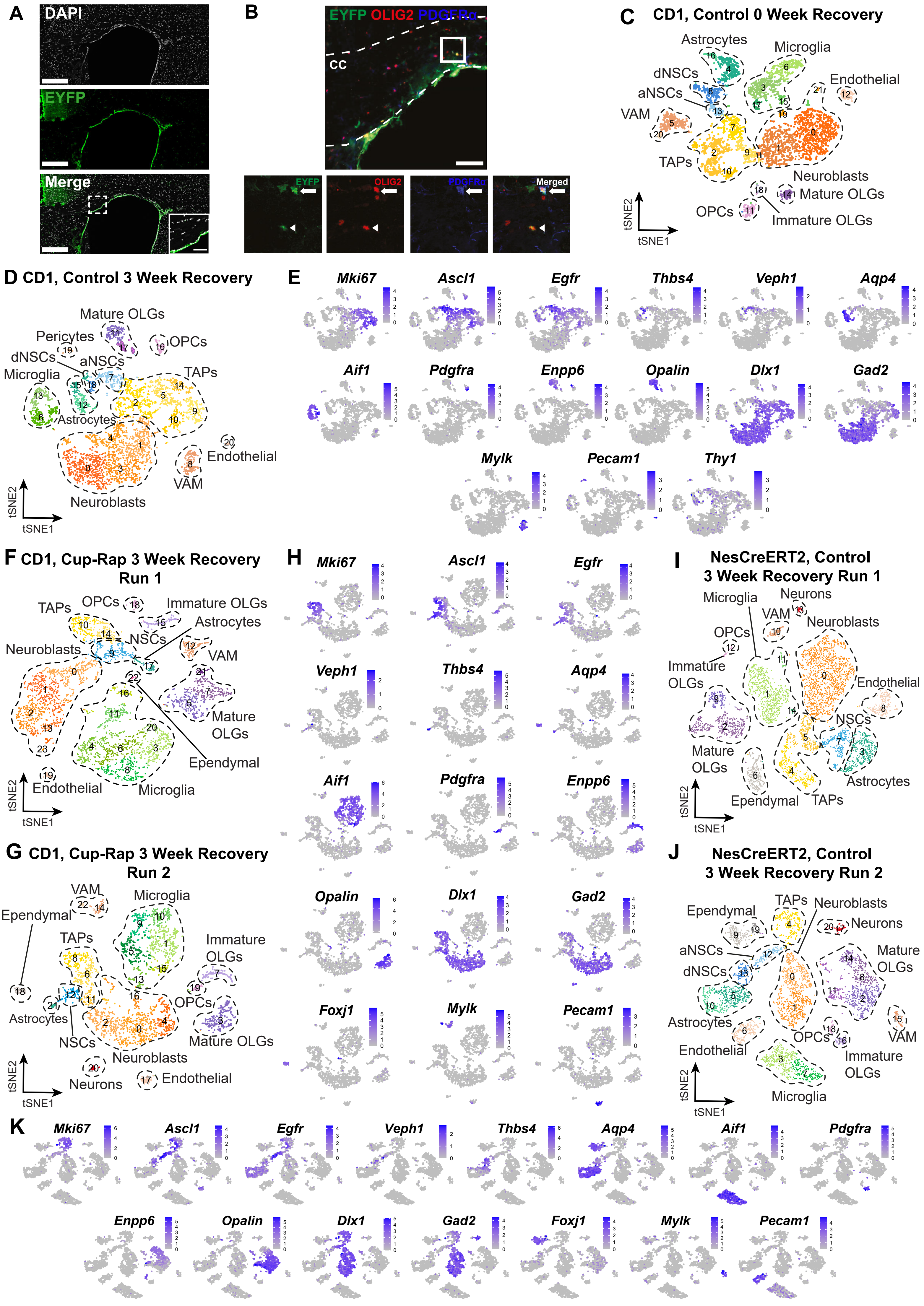

### Supplemental Figure 2

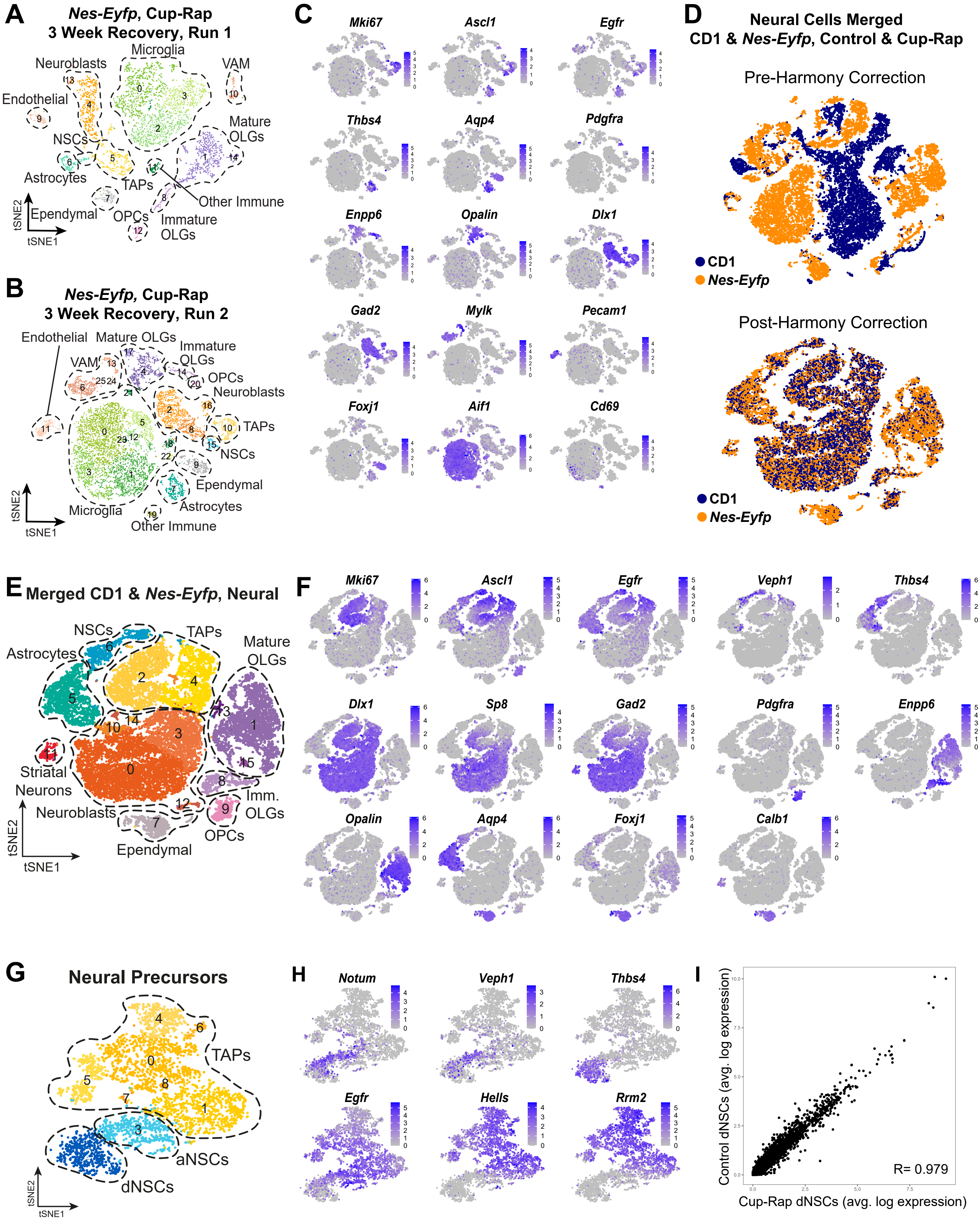

### Supplemental Figure 3

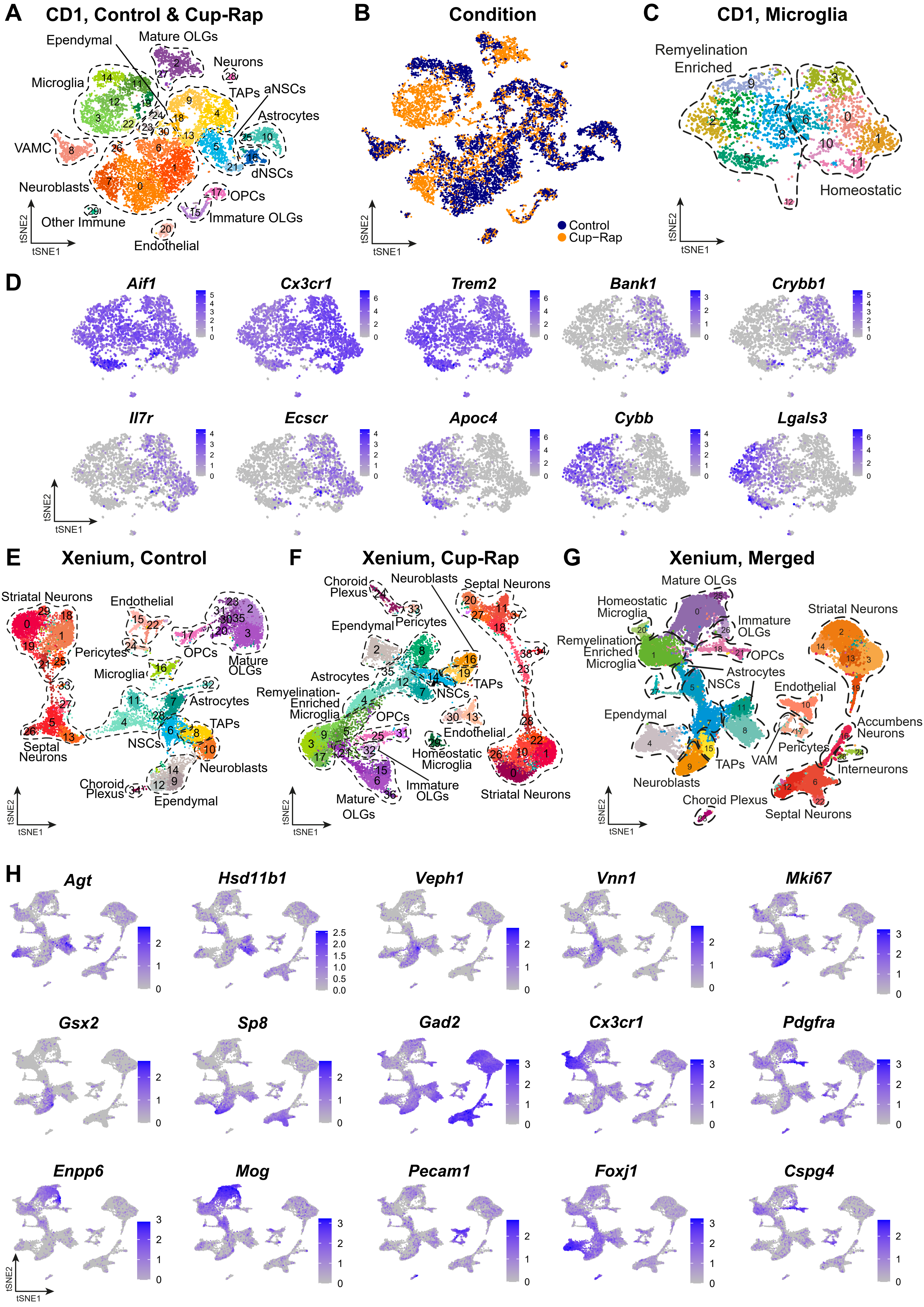

### Supplemental Figure 4

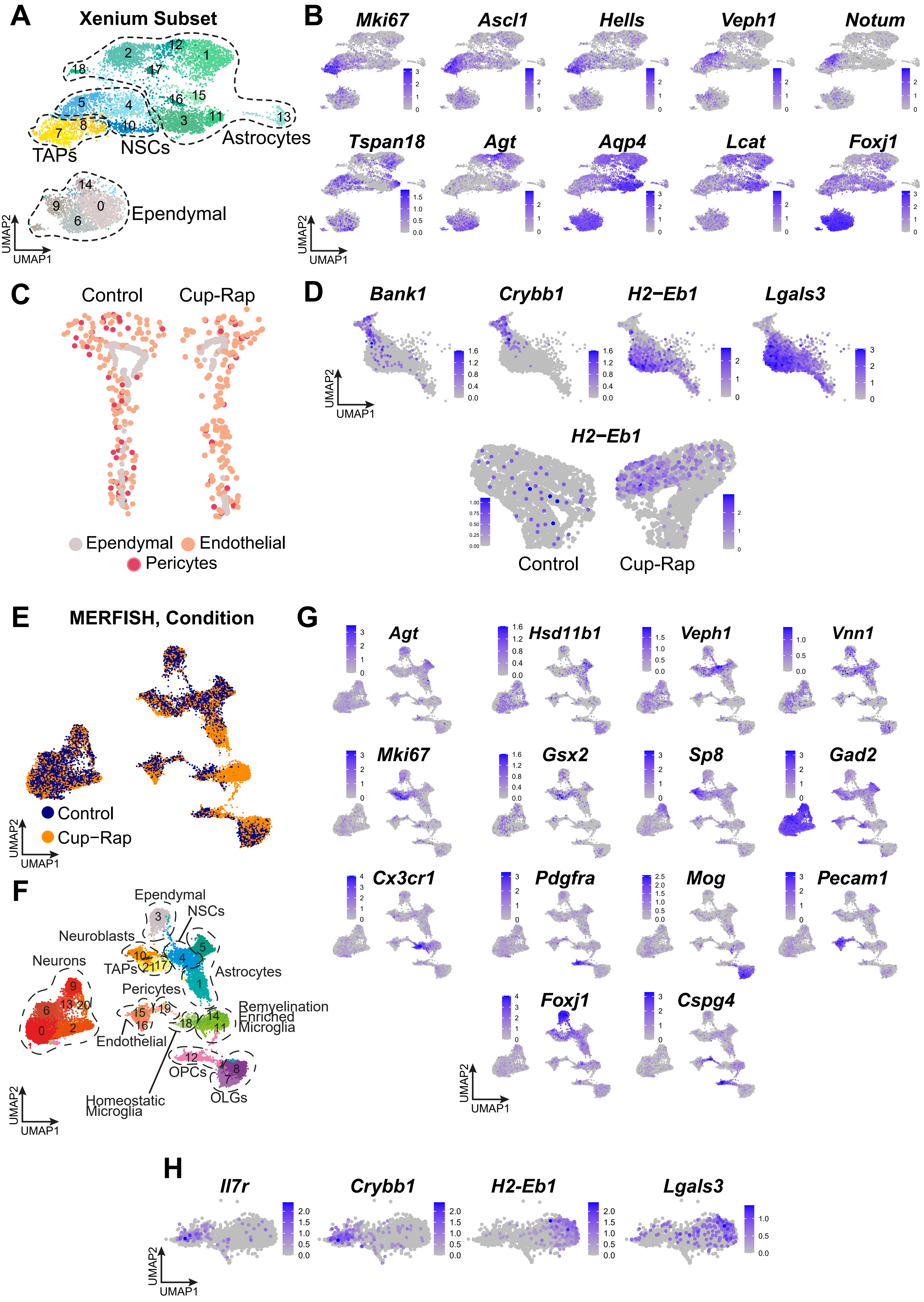
